# Integrative pan-cancer analysis suggests ISG15-linked immune and phosphoproteomic divergence between KIRC and KICH

**DOI:** 10.1101/2024.10.18.619140

**Authors:** Shrabonti Chatterjee, Joydeep Mahata

## Abstract

**Background:** Interferon-stimulated gene 15 (ISG15) is a ubiquitin-like modifier induced by inflammatory cues, yet its functional relevance in cancer remains unclear and context dependent. Although frequently used as an interferon marker, whether ISG15 reflects effective antitumor immunity or dysfunctional immune states across cancers has not been systematically resolved.

**Methods:** We performed an integrative pan-cancer analysis using TCGA, CPTAC, GEO, and immunotherapy and single-cell RNA-seq cohorts. ISG15 was examined at transcriptomic, proteomic, epigenetic, genetic, immune-infiltration, and phosphoproteomic levels. Survival-modeling, mutation and copy-number analyses, pathway-enrichment, protein–protein interaction networks, and kinase enrichment were used to define functional associations, with focused comparisons between renal clear cell carcinoma (KIRC) and chromophobe carcinoma (KICH).

**Results:** ISG15 was upregulated in all studied cancers but downregulated in KICH. Its prognostic associations were bidirectional, correlating with poor survival in KIRC, LIHC, COAD, and LGG, yet favorable outcomes in SKCM, MESO, and OV. In KIRC, high ISG15 was associated with early promoter hypomethylation, interferon signaling, immune infiltration, T-cell exhaustion signatures, and broad immune-associated phosphokinase activity. In contrast, KICH showed preserved methylation, weak interferon coupling, limited immune engagement, and phosphoproteomic enrichment of structural and trafficking pathways. Genetic alterations and deep deletions of ISG15 were associated with shorter progression-free survival.

**Conclusion:** This study indicates that ISG15 is associated with dysfunctional immune states rather than a direct effector of tumor control. These findings highlight ISG15 as a context-specific biomarker whose functional impact likely depends on tumor type and microenvironment, warranting further mechanistic and experimental validation.

## 1. Introduction

ISG15 is a member of the ubiquitin-like protein family that plays a major role in the post-translational modification of proteins, which leads to ISGylation. It is a 15 kDa protein with structural similarities, such as the presence of beta grasp folds wrapped around the central helix (ubiquitin-like proteins) [1]. ISG15 is glycosylated by three enzymes, the E1 (ISG15-activating enzyme), E2 (ISG15-conjugating), and E3 (ISG15-ligating) cascades, in which E1 activates ISG15 by binding covalently, and E2 conjugates with the active ISG15 and forms isopeptide bonds facilitated by E3 ligases [2]. During viral infection, type I interferon production is induced as part of the innate immune response. ISG15 is an important inducer of type 1 interferons, and increased ISG15 expression also upregulates the expression of conjugating and deconjugating enzymes [3]. Modulation of host and viral proteins is altered by ISGylation of target proteins such as STAT1 [4], JAK-1 [5], MDA-5 [6], Mx1, and RIG-1 [7]. Free ISG15 can be extracellular or intracellular, defining its functional aspects, as in an intracellular environment, it is conjugated with proteins that modulate or degrade it, whereas it performs immunomodulatory activity in the extracellular space [8]. Its function in immune regulation was first observed in the increased production of cytokines such as interferon-gamma from T cells and augmenting NK cell proliferation [9]. During viral or bacterial infection, it is induced by type I interferon signaling as a first line of defense [10]. Interferon expression is increased during infection, which binds to their corresponding receptors, activating downstream components such as Janus and ISF-3 (interferon-stimulating factor 3 complex), further stimulating ISGs [11]. Research has shown that ISG15 can directly influence JAK3, IRF3, and other proteins that participate in innate immunity signaling to inhibit or induce interferon-gamma secretion outside immune cells, such as monocytes, neutrophils, and lymphocytes [12]. ISG15 inhibits the ubiquitinylation of several cell proliferation/survival mediator proteins, especially kinases, which can play a role in tumorigenesis development. The exact mechanisms and pathways underlying the contribution of ISG15 to cancer progression are not well studied and need more in-depth understanding. ISG15 has a multifaceted role in cancer and can act as a tumor promoter across diverse cancers, including pancreatic ductal adenocarcinoma, nasopharyngeal carcinoma, hepatocellular carcinoma, lung cancer, colorectal cancer, endometrial carcinoma, glioma, and prostate cancer, and as a tumor suppressor in breast cancer, ovarian cancer, cervical cancer, and lung cancer. Isg15 and ISGylation dysregulation leads pathogenesis of cancer in many tissues by disrupting cytoskeletal structure, EMT, tumor microenvironment, and macrophages [13,14]. In breast cancer, IsGylation inhibits the ubiquitination and degradation of the Kirsten Rat Sarcoma Viral Oncogene Homolog (KRAS) protein, which increases cell motility and promotes Akt signaling, leading to a tumorigenic effect [15]. In ovarian cancer, it suppresses ERKI activity and increases the number of immune cells that suppress cancer stem cells [16]. Previous report demonstrates that in nasopharyngeal cancer (NPC), ISG15 promotes M2 phenotype of macrophages by activating LFA-1/SRC signaling and enhancing CCL18 production. Clinically, high ISG15+ CD163+ TAM infiltration correlates with advanced disease and poor survival, indicating a tumor-promoting role for ISG15 within the microenvironment [17]. In head and neck squamous cell carcinoma (HNSCC), it has been reported that necroptotic tumor cells release DNA that is detected by cGAS (cyclic GMP-AMP synthase) and further with help of STING signaling it activates interferons and ISG15. Further ISG15 binds to RAGE on neighboring tumor cells, that drives epithelial-mesenchymal transition (EMT) and lymphatic metastasis, helping the tumor grow and spread [18]. A recent study found that free extracellular ISG15 promotes NK cells infiltration into xenografted breast tumors in nude mice leading to suppressed breast tumor growth supporting its anti-tumor role [19]. Modulation of IL6/JAK2/STAT3 signaling by ISG15 has been reported in in one of the sub-type of renal cell carcinoma known as clear cell renal cell carcinoma (ccRCC), that directly induces renal cells proliferation [20]. In one of the rare cancer, anaplastic thyroid carcinoma (ATC), it was reported that increased ISG15 mediated ISGylation of KPNA2 leads to maintenance of cancer stem cells and decreased expression suppressed ATC growth zebrafish models [21]. Recently a pan-cancer study of ISG15 suggested strong correlation of ISG15 with immune filtration and immune check point PD-L1 in gastric and lung cancer samples. Elevated ISG15 levels lead to tumor progression and interferes tumor immune environments making them promising candidate as cancer biomarker [22]. Because of important role in tumor progression, they are also used as promising immunotherapeutic target. Recently in colorectal and renal cell carcinoma ISG15 is used as a novel tumor-associated antigen targeted by Listeria-based vaccine which increased cytokine production, and Teff/Treg ratio within the tumor microenvironment [23,24]. In different cancer types, different biological pathways governing tumor growth are altered because of the involvement of ISG15 in different pathways, but the overall presentation of ISG15 among multiple cancers is not clearly understood.

In this in silico study, we examined ISG15 expression across 22 cancer types by integrating multi-omics datasets from TCGA, GTEx, and CPTAC. We evaluated its mRNA and protein abundance and found that ISG15 is upregulated in nearly all cancers except KICH, where its expression remains low. We then assessed ISG15 within the broader context of TP53 status, mutational landscape, immune infiltration, patient survival, co-expressed genes, and pathway associations to better understand its pathogenic roles. To investigate determinants of its context-dependent immune functions, we analyzed immune modulators, stromal components, and metabolic programs linked to ISG15 activity. We also incorporated single-cell RNA-seq datasets, with a particular focus on KIRC and KICH, to resolve the cellular sources of ISG15 and identify microenvironmental cues that may drive its pro-tumor or anti-tumor behavior. Overall, this work provides an integrated view of ISG15 expression, genetic alterations, regulatory networks, and protein interactions, highlighting its complex involvement in both oncogenic signaling and antitumor responses across different tumor microenvironments.

## 2. Materials and methods

### 2.1. ISG15 expression analysis

We selected the ISG15 gene and analyzed its expression in tumor cells across different tissues utilizing patient data deposited in public repositories. We employed various databases, including GEPIA2 and UALCAN. GEPIA is a web server that extracts data from TCGA, the GTEx database of normal tissues [25], and the cBioPortal database. UALCAN is an interactive web portal for in-depth analysis of TCGA gene expression data [26]. Here, we examined ISG15 expression in GEPIA2 among all 22 types of cancer: adrenocortical carcinoma (ACC), bladder urothelial carcinoma (BLCA), breast invasive carcinoma (BRCA), cervical squamous cell carcinoma and endocervical adenocarcinoma (CESC), cholangio carcinoma (CHOL), colon adenocarcinoma (COAD), lymphoid neoplasm diffuse large B-cell lymphoma (DLBC), esophageal carcinoma (ESCA), glioblastoma multiforme (GBM), head and neck squamous cell carcinoma (HNSC), kidney chromophobe (KICH), kidney renal clear cell carcinoma (KIRC), kidney renal papillary cell carcinoma (KIRP), acute myeloid leukemia (LAML), brain lower grade glioma (LGG), liver hepatocellular carcinoma (LIHC), lung adenocarcinoma (LUAD), lung squamous cell carcinoma (LUSC), mesothelioma (MESO), ovarian serous cystadenocarcinoma (OV), pancreatic adenocarcinoma (PAAD), pheochromocytoma and paraganglioma (PCPG), prostate adenocarcinoma (PRAD), sarcoma (SARC), and skin cutaneous melanoma (SKCM). We excluded five cancer types with few control samples (n<10) or insignificant regulation (logFC<1) at the mRNA level, such as CHOL, PCPG, SARC, MESO, and UVM. We identified 22 cancer types whose log fold change (logFC)>1, with a p value <0.01 as the cutoff value. GEPIA 2 used ANOVA statistical method for differential expression on the parameter of log2(TPM+1) scale with a 0.01 q value cutoff. This program was used to explore gene regulation, as depicted by the box plot.

### 2.2. mRNA and protein expression of ISG15 across cancer stages and subtypes

We further studied the mRNA and protein expression of ISG15 in all 22 cancer types via UALCAN on the basis of pathological stages and different histological subtypes in TCGA cancer types. Clinical Proteomic Tumor Analysis Consortium (CPTAC), in UALCAN was used to study protein levels of 11 available cancer types, namely, breast cancer, ovarian cancer, lung squamous cell carcinoma, colon cancer, RCC, head and neck cancer, UCEC, lung adenocarcinoma, PAAD, glioblastoma, and hepatocellular carcinoma, via jitter plots. Among these 11 cancer types analyzed, data for ISG15 protein expression in cancer stages were not available for lung squamous cell carcinoma, glioblastoma, or hepatocellular carcinoma. Statistical changes in ISG15 mRNA and protein levels between the two stages indicate the role of ISG15 in tumor progression.

### 2.3. Survival analysis of ISG15 expression across TCGA cancer types

We conducted a survival plot analysis via the UALCAN tool for 33 TCGA cancer types to examine the impact of high and low ISG15 gene expression on patient survival in different cancer types. In addition, to understand how genetic alterations in the ISG15 gene impact patient survival across cancers, we performed progression-free survival analysis utilizing survival data from cBioPotal in patients from various cohorts. Survival plots with a p value threshold of less than 0.05 were considered significant.

### 2.4. Construction and verification of predictive nomogram

Overall survival (OS) analyses were conducted for patients with KICH and KIRC using TCGA data. Time-dependent ROC curves were generated to assess the predictive performance of ISG15 expression at 1, 2, 3, and 5 years, and AUC was calculated. Multivariate cox regression analyses were performed to identify independent prognostic factors, including age, gender, AJCC pathological stage, and ISG15 expression and Hazard ratios with 95% confidence intervals (CIs) were estimated. Variables significantly associated with OS in multivariate analysis were incorporated into nomograms to predict 1-, 2-, 3-, and 5-year OS. Nomogram performance was evaluated using calibration curves generated with 1,000 bootstrap resamples, with a 45-degree line indicating ideal agreement. Predictive accuracy was further assessed using ROC curves and corresponding AUC values. All statistical analyses were performed in R, and a two-sided p value < 0.05 was considered statistically significant.

### 2.5. TP53 mutation and methylation status of ISG15

We analyzed ISG15 expression across 22 cancer types on the basis of TP53 mutation status via the UALCAN database to determine how TP53 mutations influence ISG15 expression and progression in tumors. To assess the methylation status of ISG15, we utilized the Shiny Methylation Analysis Resource Tool (SMART). We examined the chromosomal distribution of methylation probes associated with ISG15. Additionally, CpG-aggregated methylation values against the mean expression levels in a pan-cancer context were shown with scatter plots.

### 2.6. ISG15 mutatome map mining

The cBioPortal platform was used for mutation mapping of ISG15 genes across 22 cancer types. ISG15 gene mutations, such as mutation locations, frequency, and nonsynonymous mutations across cancers were identified. Additionally, it facilitates insights into genomic alterations such as fusion, nonsense, missense, and frameshift deletions or insertions.

### 2.7. Correlation and similar gene detection in pan-cancer

We conducted a comprehensive pan-cancer analysis via GEPIA2 to identify genes that are similar to ISG15. Furthermore, we employed UALCAN, which utilizes correlation data from the TCGA database, to identify both positively and negatively correlated cancer-specific genes linked to ISG15. The Pearson correlation coefficient (PCC), indicates how the expression of one gene is related to that of ISG15. Genes with strong correlations have a PCC value exceeding 0.6, those with moderate correlations have a PCC value greater than 0.4, and genes with weak correlations have a PCC value less than 0.4.

### 2.9. Analysis of immune landscape

To investigate the relationship between ISG15 expression and specific immune cell populations, TIMER2.0 web server (http://timer.cistrome.org/) was utilized. It integrates multiple algorithms including TIMER, CIBERSORT, quanTIseq, xCell, MCP-counter, and EPIC to estimate immune cell infiltration. By utilising this server associations between ISG15 expression and natural killer T (NKT) cells, regulatory T (Treg) cells, macrophages, cancer-associated fibroblasts (CAFs), endothelial cells, common lymphoid progenitors, and myeloid-derived suppressor cells (MDSCs) were established. Additionally, TISIDB web portal (http://cis.hku.hk/TISIDB/) was employed to analyze correlations between ISG15 expression and different immune-related factors, such as tumor-infiltrating lymphocytes (TILs), immunoinhibitors, immunostimulators, major histocompatibility complex (MHC) molecules, chemokines, and their receptors.

### 2.10. Role of ISG15 in immunotherapy cohorts

To investigate the role of ISG15 in tumor immune evasion, Tumor Immune Dysfunction and Exclusion (TIDE) (http://tide.dfci.harvard.edu/login/) web platform was used. “Biomarker Evaluation” module of TIDE compared ISG15’s efficacy as a predictive biomarker against established markers across various immune checkpoint blockade (ICB) therapy cohorts. Regulator Prioritization module of TIDE was employed to evaluate the potential of ISG15 as a therapeutic target. To identify immune cell infiltration and gene expression of ISG15 across various treatment conditions both in vivo and in vitro, Gene module of Tumor Immune Syngeneic Mouse (TIMSO) database (http://tismo.cistrome.org/) was used.

### 2.11. Integrated single-ell analysis of ISG15 expression across tumor types

Single-cell transcriptomic analyses were conducted using the Tumor Immune Single-cell Hub 2 (TISCH2). TISCH2-provided UMAP embeddings were used to visualize cellular heterogeneity, with cells annotated by major lineage (including B cells, CD4□ and CD8□T cells, exhausted CD8□ T cells, dendritic cells, NK cells, monocytes/macrophages, fibroblasts, endothelial cells, and plasma cells) and by malignancy status (immune, stromal, malignant). Single-cell type datasets were used to study ISG15-associated transcriptional programs across immune, stromal, and malignant compartments. To further contextualize ISG15 function, associations with immune functional states, including interferon response, cytokine signaling, exhaustion, metabolic activity, and chromatin accessibility, were evaluated using correlation-based analyses. In parallel, pathway-restricted analyses were performed using TCGA bulk RNA-seq data for KICH and KIRC by intersecting ISG15-correlated genes with curated Hallmark gene sets from MSigDB, including interferon-gamma response, interferon-alpha response, IL2_STAT5_signaling, and IL6_JAK_STAT3_signaling. Pearson correlation distributions were compared between cancer types using the Mann– Whitney rank-sum test in GraphPad Prism. Bulk mRNA expression patterns of ISG15-associated Hallmark genes were examined using GEPIA3 to provide complementary tumor-level transcriptional context.

### 2.12. Immunohistochemistry

Immunohistochemistry study was performed using Cancer module in the HPA. In the cancer module all cancer types were selected and in each cancer type ISG15 expression is observed in the cancerous and non-cancerous/moderate cancerous tissue studies by analysing their staining intensity.

### 2.13. ISG15-associated phosphosite and kinase–substrate enrichment analysis in KICH and KIRC

Phosphoproteomic and proteomic analyses for clear cell renal cell carcinoma (ccRCC) were performed using the LinkedOmics database ISG15 expression was used as the query gene, and Spearman correlation analysis was conducted to identify phosphosites and proteins associated with ISG15 expression. Significant positive and negative associations were visualized using volcano plots and heatmaps. Functional enrichment analyses of ISG15-associated phosphoproteins were performed using LinkedOmics’ integrated Web based Gene Set Analysis Toolkit (https://www.webgestalt.org/). For non–clear cell renal cancers, publicly available phosphoproteomic datasets were obtained directly from the CPTAC. Raw phosphosite intensity data were downloaded and preprocessed independently. Kinase activity was inferred using K3Kinase enrichment analysis version 3 (https://maayanlab.cloud/kea3/#results) by assessing the enrichment of known kinase substrates for ISG15-associated phosphosites.

### 2.14. Protein □ protein interaction (PPI) and hub gene analysis

In this study, we selected genes highly correlated with ISG15 (PCC > 0.6) and moderately correlated genes (PCC > 0.4), along with 11 known tumor suppressor genes, to construct a protein □ protein interaction (PPI) network via STRING version 12.0 [27]. We then performed a cluster analysis of the network via Markov clustering (MCL) in the string. We subsequently employed the Cytohubba [28] plugin within the Cytoscape [29] platform to identify key hub genes. The top 10 hub genes were ranked on the basis of the maximum clique centrality (MCC) method within the PPI network.

### 2.15. Drug-hub gene interactions in DGIdb

We used the therapeutic-gene interaction database to detect putative drugs that target the hub genes to determine the likelihood of these genes being potential therapeutic targets. We also highlighted the drugs (approved and nonapproved) that could interact with the prognostic hub genes via DGIdb v.5.0.7 [30]. The interactions between the drug□gene network and gene categories were then visualized via Cytoscape version 3.10.1 to construct the network.

### 2.16. Pathway enrichment analysis of similar genes and hub genes

Using the clusterProfiler package in R, we performed a pathway enrichment analysis of the genes showing high and moderate correlations with ISG15. We examined similar genes that were significantly correlated (PCC > 0.4) with ISG15 to determine their gene ontology (GO) terms. GO terms with a p value less than 0.01 were considered statistically significant. Additionally, we employed ClueGO [31], a Cytoscape plug-in, to conduct a more in-depth functional analysis of the identified hub genes. Significance threshold of p ≤ 0.05 was used for pathway analysis. A kappa score threshold of 0.4 was applied. For statistical validation, the enrichment/depletion method was employed via a two-sided hypergeometric test.

## 3. Results

### 3.1. ISG15 is upregulated across multiple TCGA cancer types

Differential expression analysis revealed that the ISG15 gene was upregulated in 22 TCGA cancer types-ACC, BLCA, BRCA, CESC, COAD, BLCA, ESCA, GBM, HNSC, KIRC, LIHC, LUAD, OV, PAAD, SKCM, STAD, TGCT, THCA, THYM, UCEC, and UCS, and downregulated in KICH. The gene expression profile across all tumor samples and paired normal tissues is represented with the help of a dot plot generated in GEPIA2 at the Log2 (TPM + 1) scale. (**Figure 1a**). Additionally, individual ISG15 expression in each cancer was defined with the help of a box plot generated in GEPIA2 (**Supplementary Figure 1**). The median expression of the tumor and normal samples in different cancer was further presented in the body map across organs obtained from GEPIA2 based on TCGA and GTEx datasets. (**Figure 1b**). UCEC and HNSC had the highest mean expression values of the ISG15 gene across cancers.

**Fig. 1.**
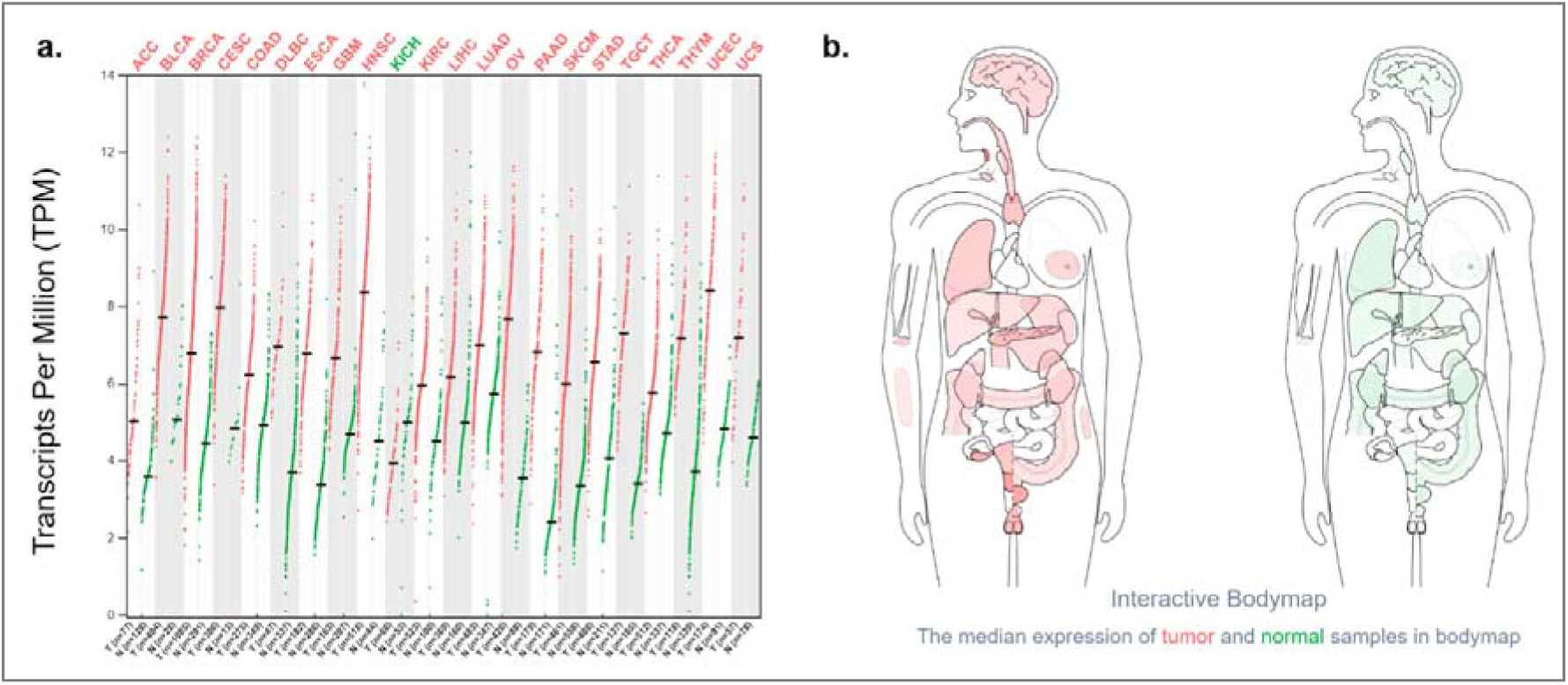
ISG15 expression in pan-cancer context. **(a)** The dot plot represents the differential gene expression level of ISG15 across 22 TCGA tumors (red dots) compared with normal samples (green dots). Y-axis represents the number of transcripts per million (log2(TPM + 1)) and X-axis denotes the number of tumor and normal samples in different cancer. **(b)** Human body map showing the median expression distribution of ISG15 in tumor and normal tissues across organs, generated using GEPIA2 based on TCGA and GTEx datasets.

### 3.2. ISG15 mRNA and protein is upregulated in cancer stages and subtypes

Next, we studied the expression level of ISG15 in individual cancer stages across 22 TCGA cancer types on the basis of patients’ pathological stage data. Our analysis revealed a significant impact of ISG15 in different stages of 13 cancer types (BLCA, HNSC, BRCA, KICH, CESC, KIRC, COAD, LUAD, ESCA, SKCM, THCA, STAD, and UCEC) among all 22 TCGA tumor types. In BLCA and ESCA, there is significant upregulation of ISG15 in stages 2, 3, and 4 compared with normal. In BRCA, there is increased expression of ISG15 in all four stages, but in stages 3 and 4, it is elevated compared with stage 1. Compared with normal samples, HNSC, CESC, and LUAD samples presented increased expression of ISG15 in all four stages. Furthermore, in UCEC and KIRC cancer types, ISG15 is markedly overexpressed in the late and early pathological stages. In KIRC stages 3 and 4, there was increased expression in stage 1 and stage 2, whereas in UCEC stages 3 and 4, there was increased expression compared with that in stage 2. In COAD, there was no significant variation in ISG15 regulation across cancer stages (**Figure 2 a-l**). ISG15 was differentially regulated in different histological subtypes of nine TCGA cancer types. In LGG, ISG15 was significantly higher in astrocytoma with respect to oligodendroglioma. In STAD, ISG15 overexpression was significant in NOS, diffuse and tubular adenocarcinoma. In BLCA, both the papillary and nonpapillary expression of ISG15 is significantly high compared to normal type. In THCA, in the classical and tall thyroid papillary carcinoma subtypes, ISG15 is significantly overexpressed. In the case of BRCA, ISG15 gene expression is increased in invasive ductal carcinoma (IDC), invasive lobular carcinoma (ILC), mixed, other, and mucinous subtypes, with the highest expression observed in the IDC subtype. In LUAD, ESCA and CESC significant ISG15 was upregulated in all the subtypes, compared to normal. COAD exhibited higher expression than normal but in subtypes significant expression of ISG15 was not there (**Supplementary Figure 2a-j**).

**Fig. 2.**
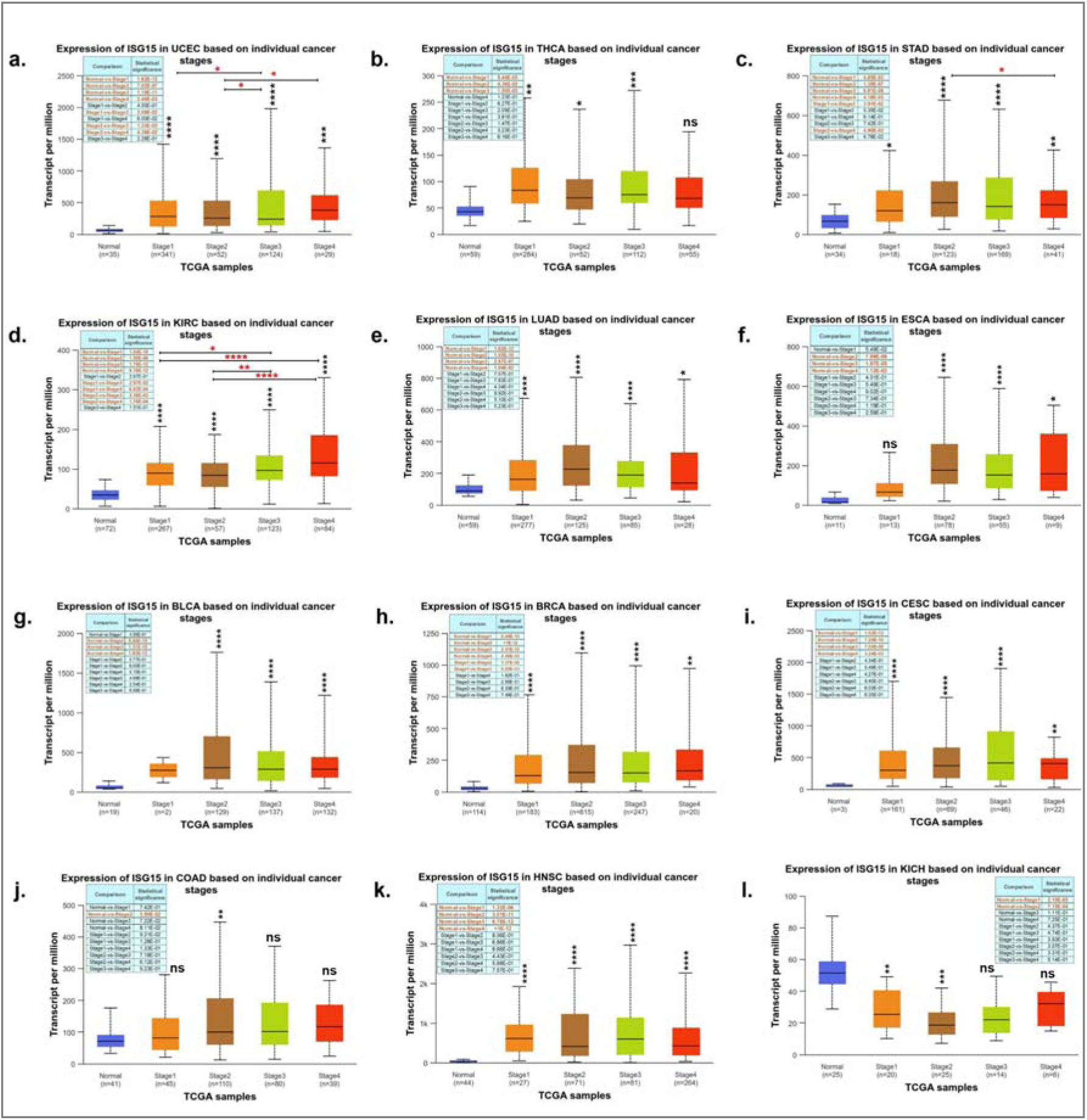
ISG15 mRNA expression landscape across TCGA tumors. **(a-l)** Box plot depicting ISG15 expression in different cancer stages. X-axis denotes the number of samples in different cancer stages. Y-axis denotes number of transcripts per million in different samples.

The expression level of the ISG15 protein was significantly high (p value <0.05) in all 10 cancer types, except for Hepatocellular carcinoma as seen in pan-cancer view (**Supplementary Figure 3a**). Further, ISG15 protein expression in each cancer type comparing tumor and matched normal samples was obtained from CPTAC portal **(Supplementary Figure 3b-l**). Among these, KIRC presented the highest level of ISG15 protein expression in tumor tissue relative to normal tissue. Hepatocellular carcinoma associated with elevated mRNA levels but suppressed protein levels suggests that the ISG15 protein might undergo rapid degradation due to the deregulation of enzymes controlling the ISGylation of ISG15 and other proteolytic systems. The protein expression levels across different cancer stages revealed increase in ISG15 protein during the later stages (stages 3 and 4) across all eight cancer types analyzed (**Supplementary Figure 4a-h**).

### 3.4. Methylation status

ISG15 methylation revealed the chromosomal distribution of 43 methylation probes associated with ISG15 that suggests it’s position within a gene-dense region transcriptionally active chromosomal segment **(Supplementary Figure 5)**. The protein-coding transcripts of ISG15 showed exon–intron organization and untranslated regions. Significant hypomethylation of the ISG15 locus in tumors compared with normal tissues is observed in BLCA, BRCA, CESC, CHOL, COAD, DLBC, ESCA, GBM, HNSC, KIRC, KIRP, LIHC, LUAD, LUSC, PAAD, PRAD, READ, SARC, SKCM, STAD, THCA, indicating epigenetic depression in many hematological malignancies. In contrast, no statistically significant difference in ISG15 methylation between tumor and normal samples is detected in ACC, DLBC, KICH, LAML, LGG, MESO, OV, TGCT, UCS and UVM, suggesting that epigenetic regulation of ISG15 is largely preserved in these cancers.

In KIRC, ISG15 promoter significant hypomethylation occurred in early-stage tumors compared to normal tissue, while later stages remain relatively unchanged. In contrast, KICH exhibits minor stage-specific differences, with only stage 3 showing a slight increase in methylation. Survival analyses indicate that in KIRC, methylation at most ISG15 CpG sites does not strongly stratify patient outcomes, consistent with subtle methylation changes, whereas in KICH, higher methylation at one CpG site correlates with poorer survival, hinting a possible prognostic role (**Supplementary Figure 6**).

### 3.5. Survival plot

Kaplan-Meier survival plot analysis revealed in SKCM, MESO and OV, patients with low/medium ISG15 expression presented a poorer survival rate than did those with high ISG15 expression. The same trend was observed at the median survival time point (defined as the point where the survival curve intersects with the 50% survival probability). COAD, KIRC, LGG, and LIHC had relatively low survival rates, which was correlated with high expression of ISG15. At the median survival time, COAD and LIHC were less different in terms of high and low/medium gene expression but were comparatively more prominent in terms of survival probability. KIRC showed a notable distinction between high and low/medium expression and their corresponding patient survival rates. In LIHC, high expression of ISG15 was almost always associated with a poor survival rate **(Figure 3a-h)**. We did not find an expression pattern in the TCGA or GEPIA2 datasets. Despite this, we included MESO in our analysis because of the observed correlations between low/medium ISG15 expression and a reduced patient survival rate. Similarly, the expression of ISG15 mRNA in LGG was not significant (logFC ∼ 0.66); however, its expression was determined to have a significant effect on patient survival rates. We found that high expression of ISG15 was associated with a lower patient survival rate.

**Fig. 3.**
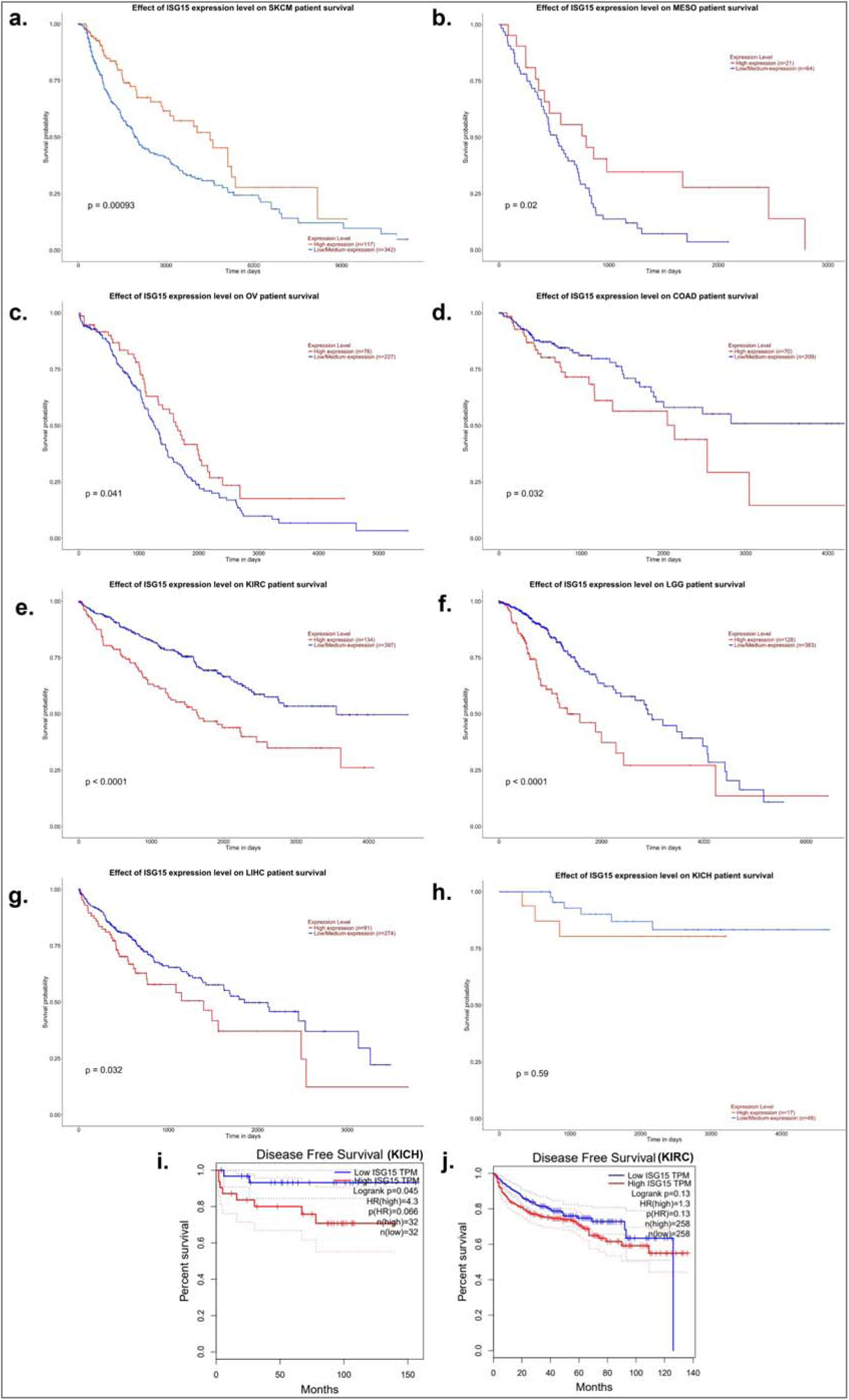
Survival analysis of ISG15 expression across cancers **(a–h)** Kaplan–Meier analyses showing improved survival with high ISG15 in SKCM, MESO, and OV, and reduced survival in COAD, LIHC, LGG, and KIRC, with no significant association in KICH**. (i, j)** Disease-free survival analyses for KICH and KIRC.

Kaplan–Meier disease-free survival (DFS) analysis revealed a differential prognostic impact of ISG15 expression across KICH and KIRC. In KICH, patients with high ISG15 expression exhibited significantly poorer DFS compared to those with low expression. This was indicated by a clear separation of survival curves (log-rank p = 0.045) and a markedly elevated hazard ratio (HR = 4.3). In contrast, patients with high ISG15 expression showed a trend toward reduced DFS, although the difference was not statistically significant (log-rank p = 0.13; HR = 1.3) in KIRC. These findings indicate that ISG15 may probably serve as a subtype-specific prognostic marker (**Figure 3 i-j**).

### 3.6. Prognostic modeling and clinical relevance of ISG15 in KICH and KIRC

To further assess the prognostic performance of ISG15, time-dependent ROC analyses were performed. The models exhibited strong short-term predictive accuracy in KICH, with a high 1-year AUC, and moderate but consistent predictive performance in KIRC across 1-, 2-, and 5-year time points, suggesting stable long-term prognostic utility **(Figure 4a, e)**. Multivariate Cox regression analysis confirmed that ISG15 expression remained an independent prognostic factor after adjustment for age, gender, and pathological stage **(Figure 4b, f)**. Based on these variables, nomograms were constructed to estimate individualized survival probabilities at 1, 2, 3, and 5 years. In the nomogram model, ISG15 contributed substantially to the total risk score, indicating its independent and clinically relevant impact on patient prognosis. Time-dependent ROC analysis demonstrated that the nomogram exhibited reliable discriminatory ability, with higher predictive accuracy in KICH and consistent performance across time points in KIRC **(Figure 4c, g)**. Calibration plots demonstrated good agreement between predicted and observed survival outcomes, indicating reliable model performance. In KICH, the close alignment of predicted and observed survival at all evaluated time points indicates good model calibration, although this should be interpreted cautiously due to limited sample size. In KIRC, calibration curves show good agreement at 1-and 3-year survival with slight deviations at 5 years **(Figure 4d, h)**.

**Fig. 4.**
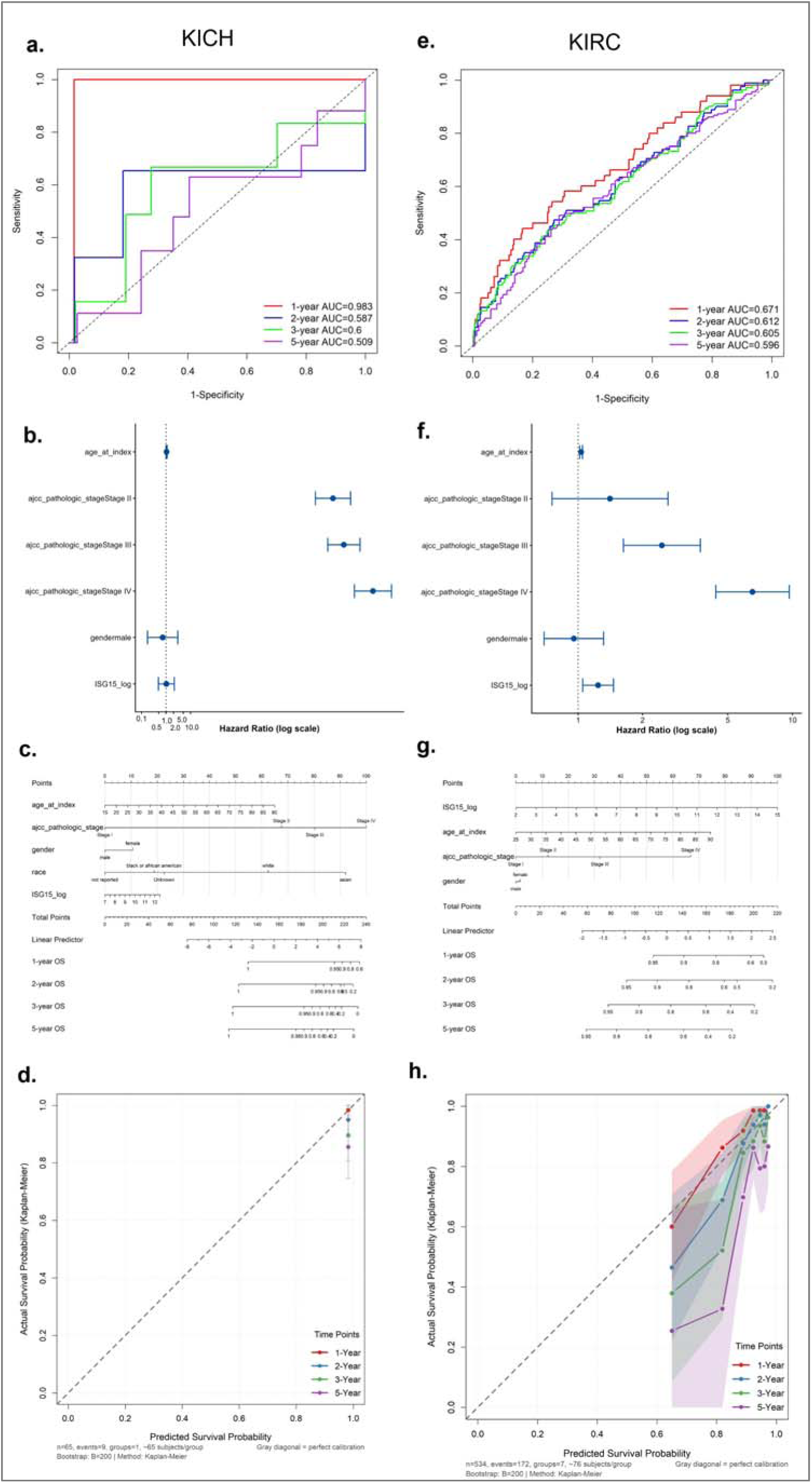
Prognostic model performance and validation in KICH and KIRC cohorts. **(a, e)** Time-dependent ROC curves for overall OS prediction. **(b, f)** Multivariate Cox regression analyses. **(c, g)** Nomogram-based survival models. **(d, h)** Calibration plots evaluating predictive performance.

### 3.7. TP53 mutation status, mutatome map, genetic alterations and PFS of ISG15

Examining ISG15 expression in relation to the TP53 mutation status provided insight into how TP53 mutations affected cancer progression **(Supplementary Figure 7)**. According to findings in breast cancer, mutations in TP53 aid in tumor progression (30). Our analysis revealed that the expression of ISG15 in TP53-mutant samples was significantly upregulated in BLCA, STAD, ESCA, READ, GBM, UCEC, LUAD, PRAD, HNSC, COAD, BRCA, and PAAD, whereas in KICH, it was downregulated with respect to its normal expression. In BLCA, UCEC, LUAD, and HNSC samples, ISG15 expression was significantly elevated in both TP53-mutant and nonmutant samples compared with normal samples. In KICH, ISG15 expression is significantly lower in both TP53 mutant and nonmutant samples than in normal tissue, with the lowest expression in nonmutant samples.

ISG15 mutation study in 10,967 patient samples from the TCGA revealed that ISG15 is mutated in approximately 2% of cases, with missense mutations and a smaller fraction of samples presented truncating mutations, amplifications, and deep deletions **(Supplementary Figure 8a-b).** The most frequently observed mutation types were missense mutations, such as G68S in lung squamous cell carcinoma and E127K in cutaneous melanoma, with varying allele frequencies. Gene copy alterations included gain of function in two cases (lung squamous cell carcinoma and cutaneous melanoma) and deletions in cervical squamous cell carcinoma and invasive ductal carcinoma of the breast. The presence of nonsense mutations, such as E160 and E27 in colorectal adenocarcinoma and cervical squamous cell carcinoma, respectively, indicates potential loss-of-function events **(Supplementary Figure 8c)**. We observed that amplification and deep deletion are the most prevalent genetic alterations found in ISG15 gene modifications across all cancer types **(Supplementary Figure 8d)**.

The PFS results, with a significant p value of 0.0173, revealed that alterations in ISG15 may be associated with a poor prognosis. The PFS predicts the percentage of patients whose disease has not progressed or worsened over time. The altered group in PFS declined more sharply in patients with ISG15 gene mutations, suggesting a shorter time until disease progression. **(Supplementary Figure 9a)**.

mRNA versus copy-number alterations (CANs) reflected decreased ISG15 expression due to deep deletions and increased expression due to amplification. Shallow deletions and diploid samples didn’t affect much suggesting that a single copy loss does not significantly impact ISG15 expression **(Supplementary Figure 9b)**.

### 3.8. Role of ISG15 in immunotherapy cohorts

We performed a comparative analysis to evaluate the biomarker potential of ISG15 in relation to previously reported biomarkers. In melanoma cohorts, particularly those treated with anti– PD-1, anti–CTLA-4, or combination therapy, ISG15 showed AUC values in the range of ∼0.6–0.75. In these same cohorts, immune-related markers such as CD8, IFNG, and the Merck18 T cell–inflamed signature revealed AUCs more than 0.7. In NSCLC and HNSC cohorts, ISG15 displayed AUC values around 0.6 similar to CD274 (PD-L1), IFNG, and CD8. In KIRC those treated with anti–PD-1, ISG15 demonstrated AUC values, similar to those of TMB and MSI. IFNG displayed low AUC values in this cohort, and CD8, CD274 and T. Clonality markers showed consistent predictive power. In glioblastoma cohorts, ISG15 showed low AUC values in immune activation markers - CD8, IFNG, and Merck18 except in TIDE and CD274. Biomarkers such as CD8, IFNγ, and the Merck18 T-cell-inflamed signature display higher AUC values, suggesting that gastric tumors responding to PD-1 therapy are characterized by high cytotoxic T-cell infiltration and strong interferon-γ signaling **(Supplementary Figure 10a)**. In immunotherapy cohorts treated with anti-PD-1, anti-CTLA-4, or combination regimens, Cox proportional hazards analyses showed adverse associations between ISG15 expression and clinical outcome, suggesting that high ISG15 does not reliably predict benefit from checkpoint blockade and may instead reflect resistance driven by immune exhaustion. Immune cell–type–resolved expression datasets demonstrate enrichment of ISG15 in immunosuppressive compartments, including M2 macrophages, MDSCs, and cancer-associated fibroblasts, linking ISG15 expression to a suppressive tumor microenvironment CRISPR-Cas9 immune screen datasets further support a functional role, as ISG15 shows significant log2 fold-change signals, implying that modulation of ISG15 impacts immune–tumor interactions rather than acting as a passive interferon marker **(Supplementary Figure 10b)**.

Further analysis in TIMSO showed ISG15 expression is dynamically regulated in response to immune checkpoint blockade across multiple immunotherapy-treated cohorts, with clear differences between responders and non-responders. Across diverse tumor models, ISG15 is frequently upregulated following anti–PD-1, or anti–CTLA-4 therapy, and this induction is more pronounced in responders **(Supplementary Figure 11a)**. ISG15 expression in response to different inflammatory cytokine stimulations across multiple mouse tumor models revealed there is a consistent and strong induction of ISG15 in mentioned tumor types, with IFNγ producing the largest increase. The statistical annotations indicate significant increase compared with baseline in most models. In melanoma, ISG15 exhibited elevated expression and positive Z-scores across multiple PD-1 and CTLA-4 treated cohorts **(Supplementary Figure 11b)**. High ISG15 levels were associated with increased T-cell dysfunction signatures. Collectively, these datasets converge on the observation that ISG15 marks chronic interferon signaling associated with immune dysfunction, suppression, and variable immunotherapy responsiveness across cancers. ISG15 shows relative enrichment in specific tumor microenvironment components, including M2-like tumor-associated macrophages (TAM M2) and FAP-positive cancer-associated fibroblasts (CAF FAP), with high normalized Z-scores in these cell-type–specific datasets.

### 3.9. Immune infiltration, immunomodulatory associations, and immunohistochemical characterization of ISG15

TIMER 2.0 database was used to identify immune infiltration due to ISG15 expression all the cancer types. Global immune correlation patterns demonstrated that ISG15 exhibits associations with immune infiltrates across all cancers, most prominently macrophage populations (M0, M1, and M2) as estimated by CIBERSORT, TIMER, EPIC, and MCP-counter, indicating that ISG15 is closely coupled to macrophage recruitment and polarization at a pan-cancer level **(Figure 5a)**. In KICH, ISG15 expression showed strong positive correlations with total macrophages and classically activated macrophage subsets, including macrophage EPIC (r = 0.528), macrophage XCELL (r = 0.346), and macrophage M1 signatures across CIBERSORT and CIBERSORT-ABS. M2 macrophage infiltration exhibited mixed but overall positive associations in KICH, suggesting a macrophage-rich microenvironment with active innate immune engagement. In contrast, KIRC demonstrated weaker or negative correlations with macrophage populations, particularly for M2 macrophages (e.g., macrophage M2 QUANTISEQ r =-0.337, XCELL r =-0.18). This pattern suggests a relative attenuation of macrophage recruitment or altered macrophage functional states in KIRC. In KICH, ISG15 expression correlated positively with NK cells across EPIC, MCP-COUNTER, and CIBERSORT-ABS algorithms (r values up to 0.429), indicating enhanced innate cytotoxic potential. Conversely, KIRC showed positive and negative NK cell associations in different methods. This suggests impaired NK-mediated tumor surveillance in KIRC despite elevated ISG15 expression. Analysis of T cell subsets revealed fundamentally different adaptive immune states between the two cancer types. In KICH, ISG15 expression was negatively associated with CD8□ T cell infiltration and CD4□ effector memory subsets, indicating limited adaptive immune engagement despite strong innate signaling. In contrast, KIRC exhibited significant positive correlations between ISG15 and multiple CD8□ T cell populations, including CD8□T cells estimated by TIMER (r = 0.328), CIBERSORT, and XCELL, as well as CD8□ effector and central memory subsets **(Figure 5b)**. However, this apparent T cell enrichment coincided with positive associations with exhaustion markers and immune checkpoint-related gene signatures suggesting that ISG15 in KIRC is linked to a dysfunctional or exhausted T cell phenotype rather than effective antitumor immunity.

**Fig. 5.**
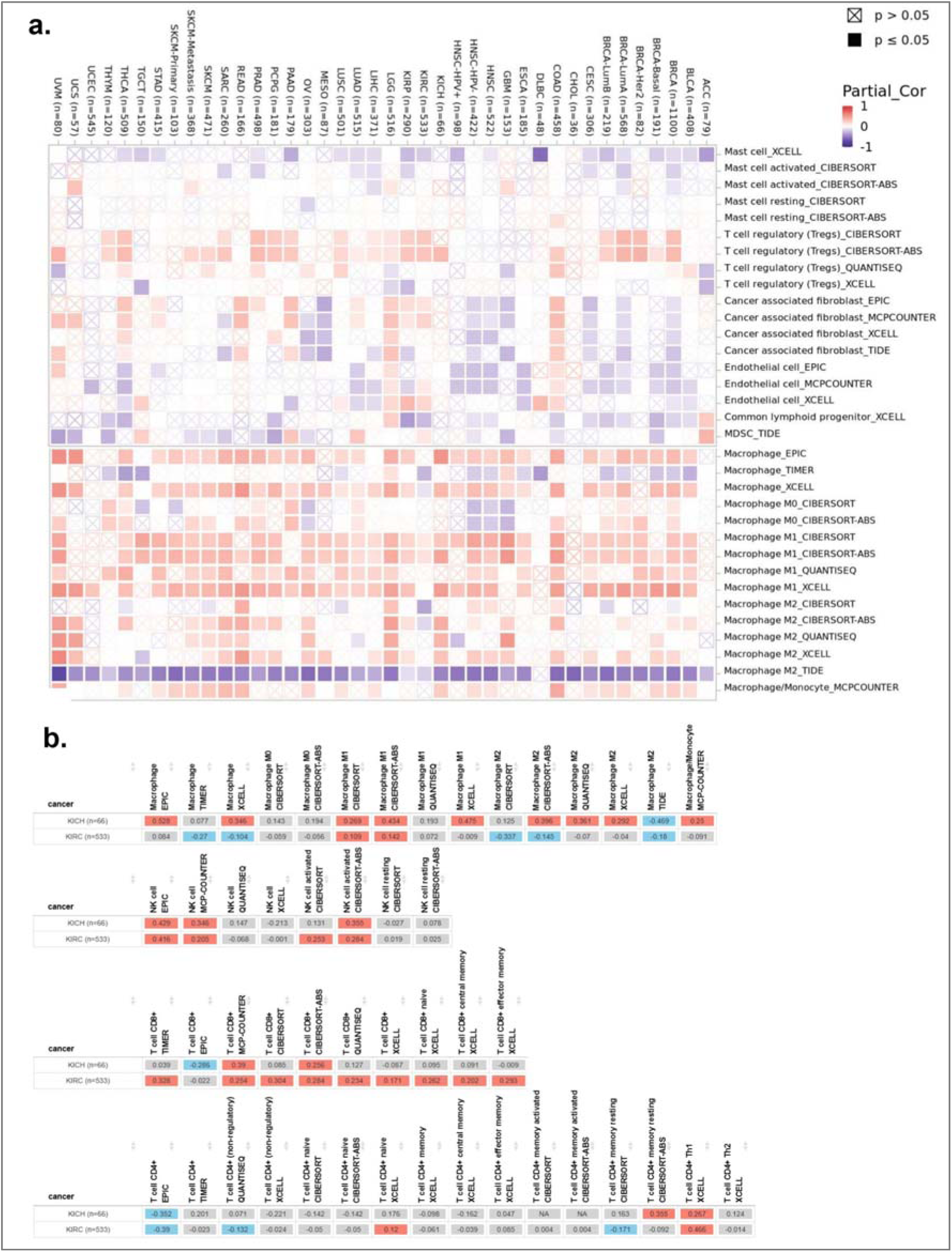
Correlation of ISG15 with immune features across cancers and renal subtypes. **(a)** Heatmap showing purity-adjusted partial Spearman correlations between ISG15 expression and immune cell infiltration across cancers. **(b)** Comparative ISG15–immune infiltration patterns in KIRC versus KICH.

Heatmap analyses of immune gene sets revealed that KICH displayed stronger associations between ISG15 and antigen presentation machinery, including MHC class I and II genes, interferon-stimulated genes, and innate immune signaling components. This supports a model in which ISG15 contributes to an interferon-responsive, immunologically active microenvironment in KICH. KIRC demonstrated enrichment of inflammatory signaling without coordinated antigen presentation **(Figure 6a-f)**.

**Fig. 6.**
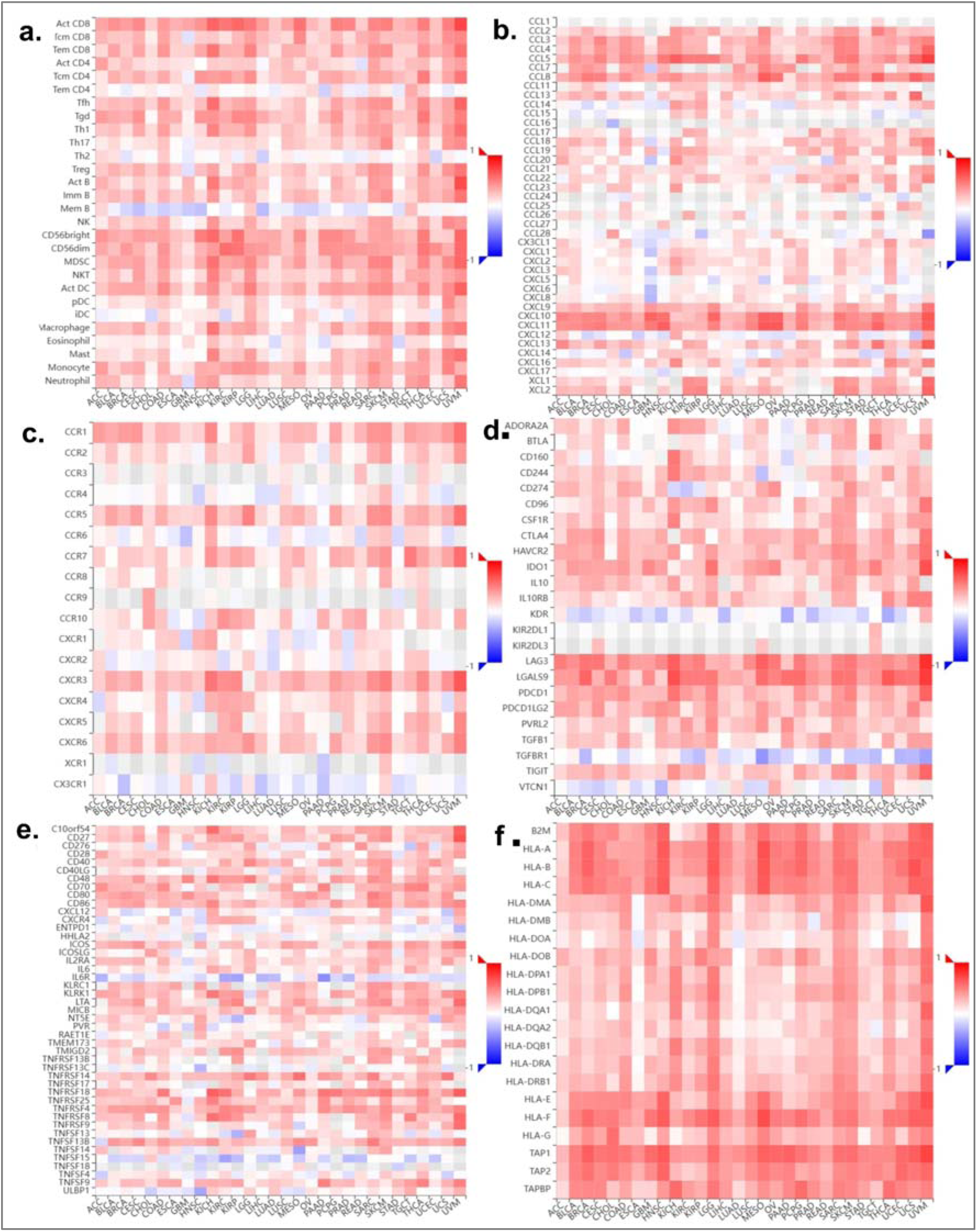
Correlation of ISG15 with tumor-infiltrating lymphocytes. **(a)** Immune cell subsets, like CD8□ T cells, CD4□ T cells, B cells, NK cells, macrophages, dendritic cells, and other immune populations, **(b)** chemokines, **(c)** chemokine receptors, **(d)** Immune checkpoint and immunoregulatory molecules (inhibitory and stimulatory markers), **(e)** Immunostimulatory genes, and **(f)** Major histocompatibility complex (MHC) molecules across cancer types. X-axis denotes cancer types, Y-axis immune components.

In immunohistochemistry of ISG15, HPA004627 antibody was used to detect the stained tissues for ISG15 expression in the cancerous tissue with respect to moderate or non-cancerous tissue in each cancer type mentioned in the HPA. This analysis revealed higher number of cases in cervical cancer, thyroid cancer, stomach cancer, pancreatic cancer and liver cancer. Immunohistochemistry images reveal predominantly moderate-to-strong cytoplasmic staining with a granular pattern, indicating that ISG15 is mainly localized in the cytoplasm of tumor cells (**Supplementary Figure 12)**.

### 3.10. Single-cell characterization revealed cell-type specific expression patterns associated with ISG15 in RCC subtype

Across multiple independent KIRC single-cell RNA-seq datasets, UMAP projections reveal a consistent segregation of malignant, immune, and stromal compartments, with ISG15 expression showing strong cell-type specificity rather than uniform induction. ISG15 signal is predominantly enriched within immune clusters, particularly myeloid populations and subsets of T cells, while malignant epithelial cells show comparatively lower and heterogeneous expression. This spatial restriction across datasets suggests that ISG15 is not merely a tumor-intrinsic interferon marker but play role in shaping immune behavior, immune suppression, or immune exhaustion within the tumor **(Figure 7a-f)**. In contrast to KIRC, the KICH single-cell dataset demonstrates markedly reduced ISG15 expression across both immune and non-immune compartments, with sparse and weak signal intensity. Across immune populations, including T cells, myeloid cells, and other leukocyte subsets, ISG15 expression was detected only in a small fraction of cells and lacked strong cluster-specific enrichment. The attenuated ISG15 signal in KICH implies fundamental differences in interferon signaling tone, immune activation, or immune infiltration between ccRCC and chromophobe RCC **(Figure 7 g)**. Combined overall expression of ISG15 across different cell types in KICH and KIRC is shown with help of heatmap **(Figure 7h).**

**Fig. 7.**
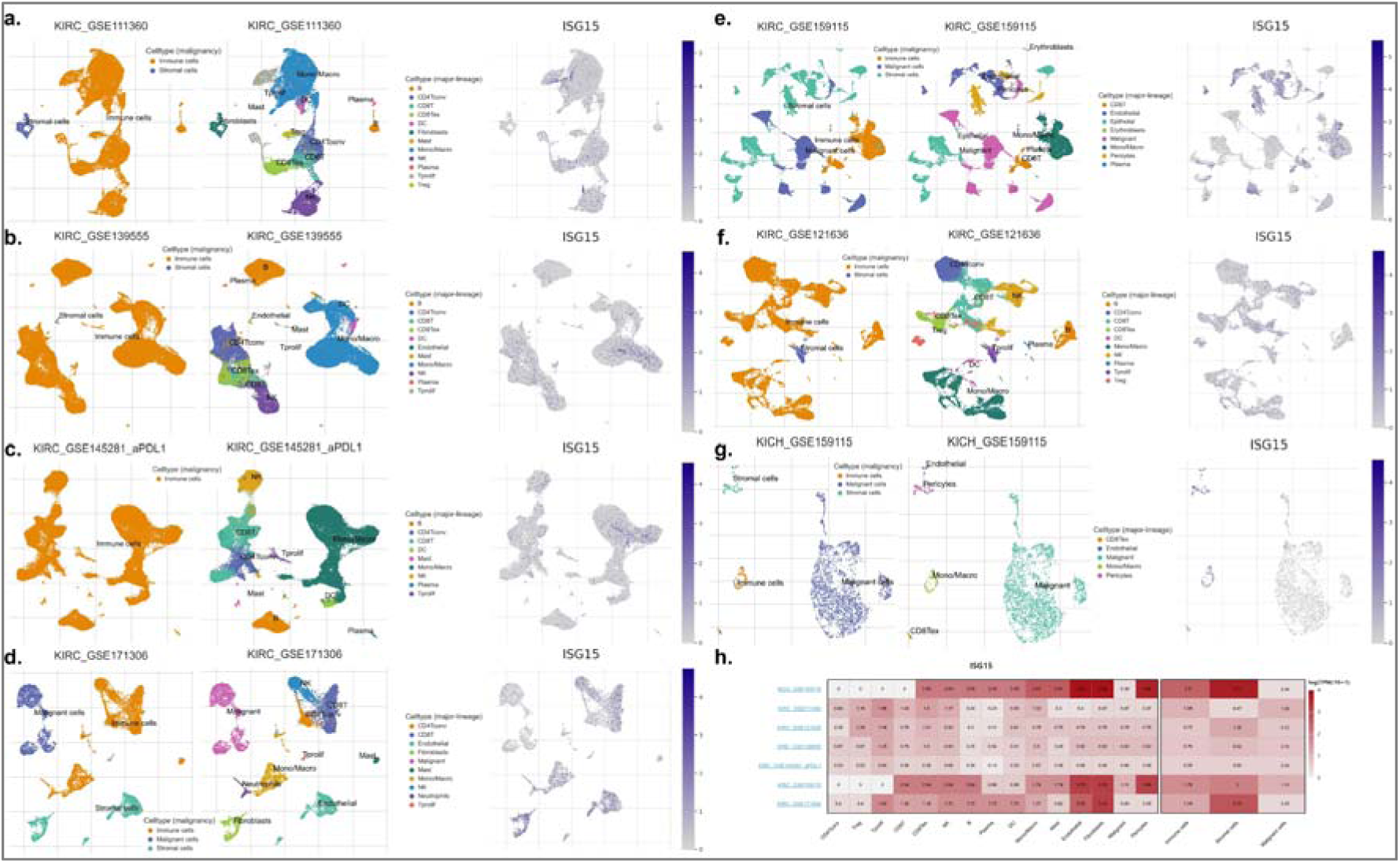
Single-cell ISG15 expression across immune, malignant, and stromal compartments in KIRC **(a–f)** and KICH **(g)**. **(h)** Heatmap of average ISG15 expression per cell type across datasets.

The correlation heatmap shows that ISG15 exhibits strong positive associations with interferon-stimulated genes, antigen presentation machinery, and immune activation markers across KIRC datasets, whereas these correlations are substantially weaker or absent in KICH. Across immune populations, including T cells, myeloid cells, and other leukocyte subsets, ISG15 expression was detected only in a small fraction of cells and lacked strong cluster-specific enrichment. In stromal compartments, including fibroblasts and endothelial cells also showed minimal ISG15 expression, with no evident spatial co-localization with immune clusters. Cell-type resolved analysis reveals that ISG15 correlations are strongest within myeloid cells and T cells, with comparatively modest or inconsistent associations in endothelial and fibroblast populations. Dataset-level concordance further indicates that ISG15-immune relationships are not driven by a single cohort or technical artifact. These findings highlight immune cell–specific integration of ISG15 into functional signaling programs, supporting a context-dependent role tied to immune cell states **(Figure 8a)**.

**Fig. 8.**
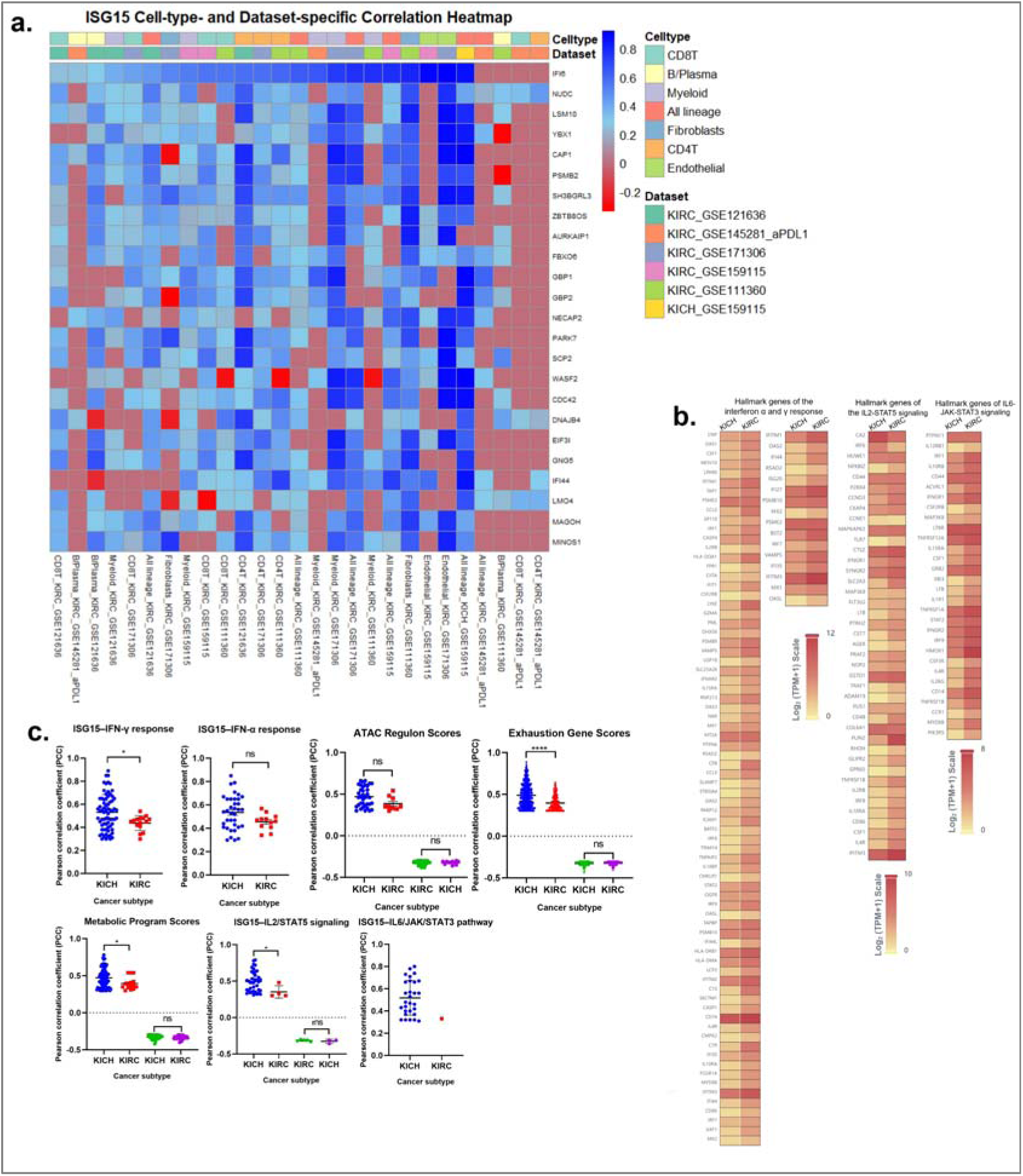
Immune cells association with ISG15 **(a)** Cell type and dataset specific Pearson correlations between ISG15 and coexpressed genes from TISCH2 **(b)** Associations between ISG15 and Hallmark immune functional signatures, including interferon response and exhaustion, in KIRC versus KICH. **(c)** Bulk tumor expression of ISG15-associated Hallmark genes in KIRC and KICH from TCGA.

In KIRC, hallmark gene analysis revealed, high ISG15 expression strongly associated with enrichment of immune and interferon-related hallmark pathways, including interferon-α response, interferon-/ response, inflammatory response, and JAK-STAT signaling. In contrast, KICH demonstrated minimal or absent enrichment of interferon and immune hallmarks in relation to ISG15 expression. Hallmark scores in KICH were generally weak, inconsistent, or uncoupled from ISG15 levels, reflecting a lower interferon signaling tone and reduced immune engagement **(Figure 8b).** Comparative analysis of immune functional scores demonstrates that higher ISG15 expression in KIRC is associated with altered T-cell exhaustion signatures, interferon-/ response, and JAK-STAT signaling, whereas these associations are non-significant in KICH. Notably, ISG15 does not uniformly correlate with increased immune fitness; instead, its association with exhaustion and metabolic programs suggests a complex role in shaping dysfunctional or chronically stimulated immune states within ccRCC **(Figure 8c)**.

### 3.11. Correlating and similar gene detection of ISG15

Correlating genes analysis revealed that 7 common genes-interferon protein 35 (IFI35), interferon-induced protein 44 (IFI44), 2’-5’-oligoadenylate synthetase like (OASL), MX dynamin-like GTPase 1 (MX1), radical S-adenosyl methionine domain containing 2 (RSAD2), 2’-5’-oligoadenylate synthetase 2 (OAS2), and interferon regulatory factor 7 (IRF7) are uniformly present across all cancer types except STAD **(Figure 9b).** We obtained another set of 31 similar genes with moderate correlations (Supporting Information Table 1). And detailed set of correlating genes (Supporting Information Table 2). The correlation analysis between ISG15 and seven commonly associated genes was analyzed based on their expression levels in TCGA tumor and normal tissues. The results revealed that, except for OAS2, the remaining six genes exhibited a mean positive correlation coefficient greater than 0.6 across all TCGA cancer types **(Figure 9a).** A multiple-gene comparison of these seven genes showed significant upregulation of all seven genes in tumor tissues of HNSC, PAAD, and TGCT. LAML, IFI44, MX1, RSAD2 and OAS2 were prominently upregulated relative to their basal expression levels. Conversely, KICH exhibited a significant downregulation **(Figure 9c)**. Further, an OS analysis of these genes revealed that in LGG, high expression of all seven genes was associated with poorer survival outcomes, whereas in SKCM, lower expression correlated with reduced survival **(Figure 9d).**

**Fig. 9.**
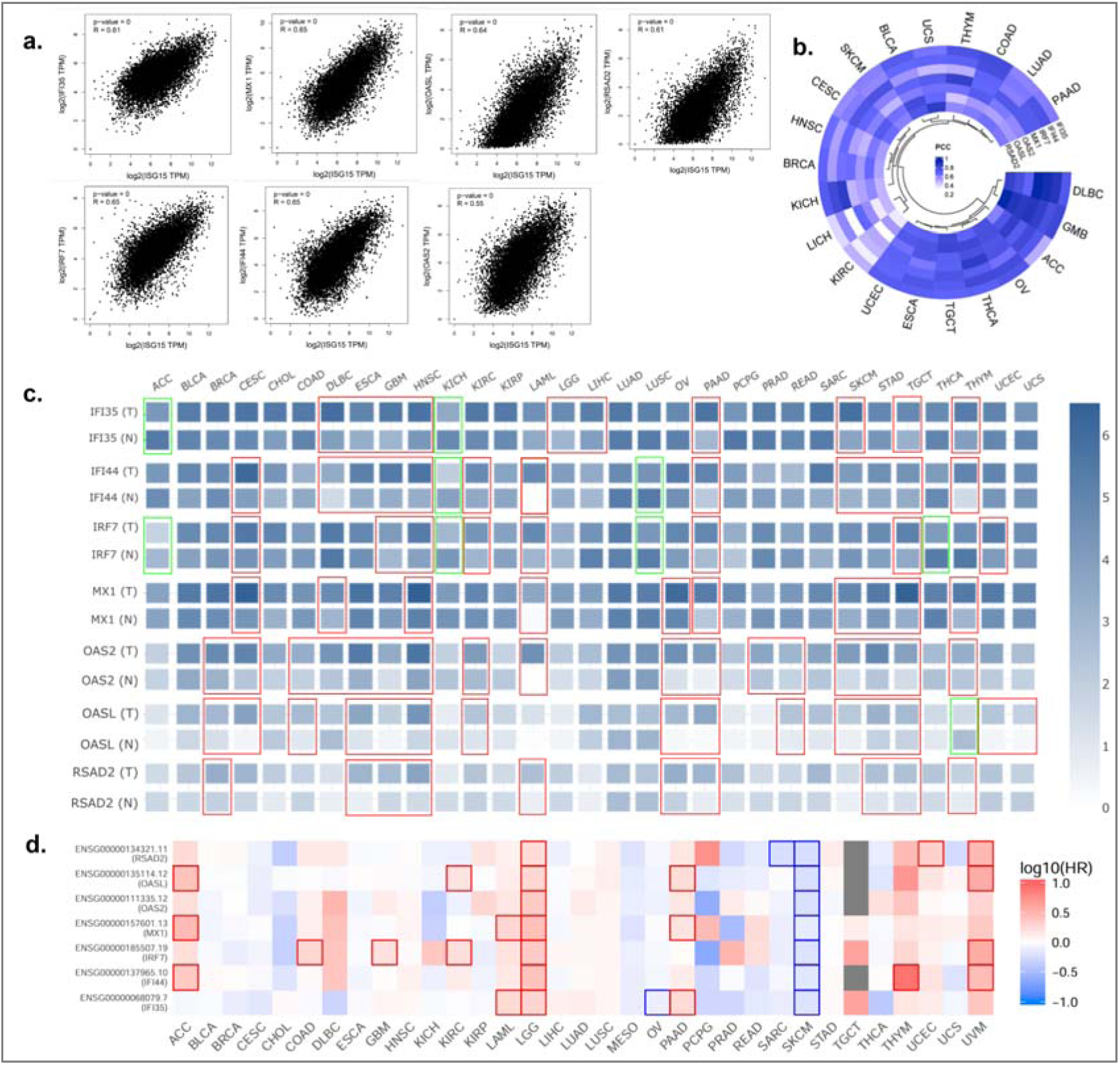
This figure summarizes the correlation, expression, and survival relevance of ISG15-associated genes across cancers. **(a, b)** Correlation patterns and Pearson correlation heatmap showing consistent positive associations between ISG15 and seven common genes across 21 cancer types. **(c)** Tissue-wise expression of these genes in tumor and normal samples across cancers, shown as log2(TPM + 1). **(d)** Overall survival associations of the seven genes across cancer types, assessed by the Mantel–Cox test, with colored borders indicating statistical significance.

### 3.12. STRING analysis of the PPI network and hub genes

After identifying genes similar to ISG15, we constructed a protein–protein interaction (PPI) network of the 39 obtained genes. Among 39 genes, 11 tumor suppressor genes (including TP53, RB1, BRCA1, BRCA2, PTEN, APC, CDKN2A, VHL, NF1, SMAD4, and WT1) and ISG15, with moderate (r > 0.4) and strong (r > 0.6) correlations, revealed a network comprising 60 nodes and 1003 edges (**Supplementary Figure 13a).** Hub genes obtained from PPI networks were ISG15, STAT1, IFIT2, OASL, IFI35, IFI44L, IFI44, IFIT1, IFIT3, and IRF7 **(Supplementary Figure 13b),** (Supporting Information Table 3). Further investigation revealed that 15 genes, IFNB1, HERC5, XAF1, DDX58, IRF7, STAT1, PSMB9, TRIM21, TYMP, FBXO6, NMI, MX1, IFI35, PSME2 and ISG15, from this PPI network interact with tumor suppressor genes (**Supplementary Figure 13c**). Drugs targeting the identified hub genes were shown in **Supplementary Figure 13d**.

### 3.13. Pathway enrichment and Gene Ontology analysis

We identified 141 significant GO terms (p value < 0.01), consisting of major pathways related to viral processes, interferons, and immune regulation (Supporting Information Table 4). Some of them are the defense response to the virus, regulation of the viral life cycle, regulation of the innate immune response, the MDA-5 signaling pathway, the cytokine-mediated signaling pathway, type I interferon production, the RIG-I signaling pathway, etc. The top 40 GO terms were plotted for visualization **(Supplementary Figure 14a)**. ClueGO analysis of the hub genes in the ISG15 network revealed three major biological processes: the cellular response to type 1 interferon, the expression of IFN-induced genes, and the expression of IFNG-stimulated genes **(Supplementary Figure 14b-d)**.

### 3.14. Phosphoproteomic pathway enrichment reveals dysregulated signaling networks in RCC

Phosphoproteome analysis revealed regulated phosphoproteomic signature influenced by ISG15 in KIRC, shown in the heatmap with both positive and negative correlations. The ISG15-correlated phosphoproteins include signaling and adaptor molecules such as MAPK1, MAPK3, AKT1, SRC, EGFR, ABL1, CHUK, TBK1, and CDK1**(Figure 10a, b)**. Gene ontology enrichment of phosphoproteome shows significant enrichment of interferon response, B-cell activation, RNA processing, DNA damage repair, and leukocyte differentiation, were consistent with an immune-and stress-responsive signaling environment **(Supplementary Figure 15a)**. Kinase enrichment analysis revealed kinases involved in SRC family kinases (SRC, LYN, FYN), MAPK pathway kinases (MAPK1/3, MAP2K1/5), AKT1, CDKs, and PRKACA, whose substrates are distributed across immune signaling, cell cycle regulation, and transcriptional control **(Figure 10c)**. In contrast, the ISG15-associated phosphoproteome in KICH were dominated by processes such as actin filament organization, protein localization to the nucleus, intracellular transport, and cytoskeletal assembly, rather than immune or interferon-related pathways (**Supplementary Figure 15b**). Kinase enrichment highlighted kinases, including MAPK14, MAPKAPK2, RPS6KA1, AKT1, and PAK1, with fewer SRC-family and immune-associated kinases **(Figure 10d)** compared with KIRC that are inclined towards structural and trafficking-associated proteins rather than immune signaling mediators. The relative absence of strong immune-related phosphosignatures in KICH aligns with its known lower immune infiltration and more epithelial, metabolically stable tumor phenotype. ISG15 expression probably reflects broader cellular states that differ fundamentally between renal cancer subtypes.

**Fig. 10.**
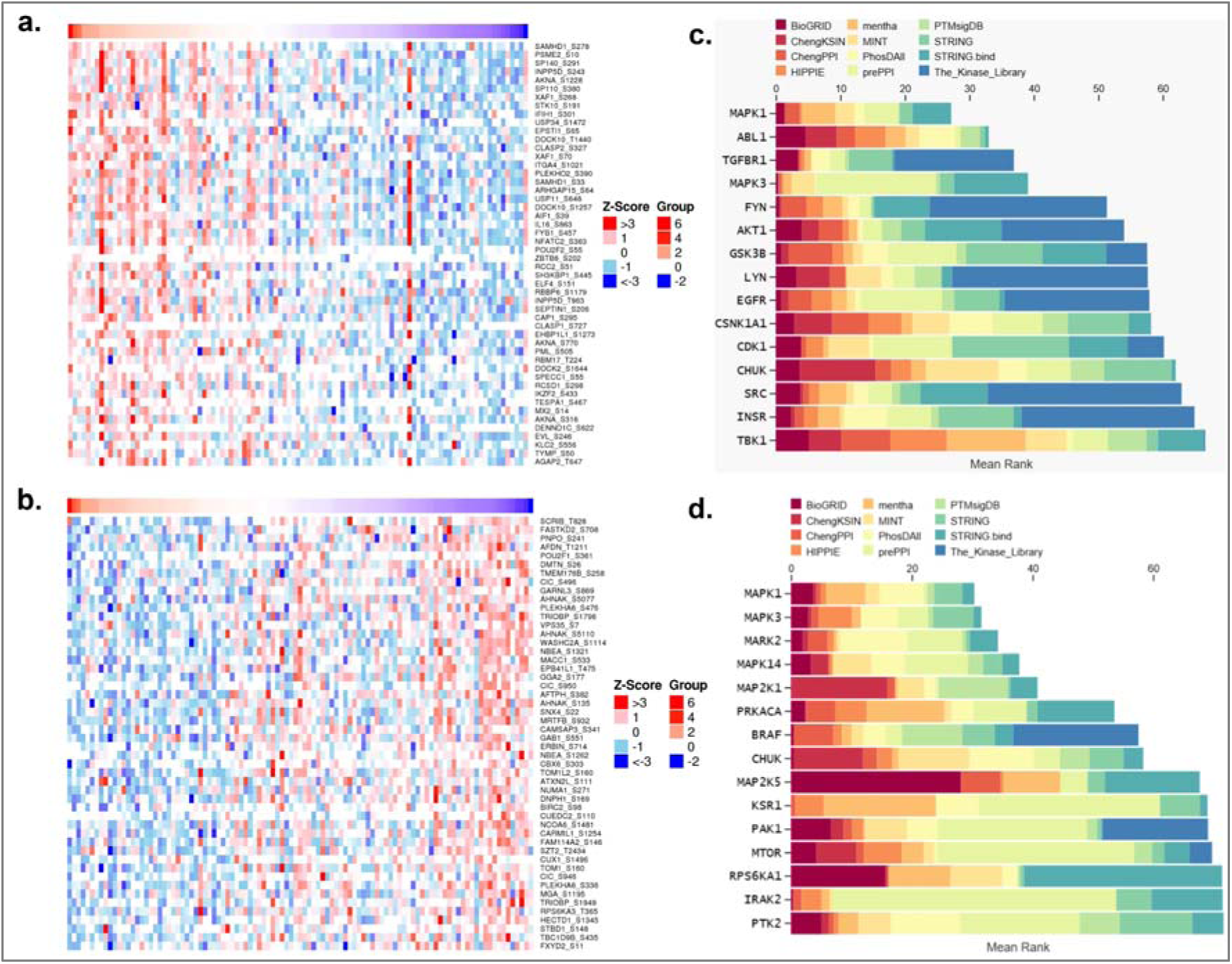
ISG15-associated phosphoproteomic and kinase signaling in renal cancer subtypes. **(a, b)** Heatmaps of phosphoproteins positively and negatively correlated with ISG15 in KIRC (Z-score normalized). **(c, d)** Kinase enrichment analysis of ISG15-associated phosphoproteins in KIRC, and KICH respectively.

## 4. Discussion and Conclusion

ISG15 shows a context dependent prognostic role, associating with poor outcomes in liver, brain, colorectal, and renal cancers, but favorable survival in ovarian, mesothelioma, and skin cancers, highlighting its dual antitumor and protumor potential [24]. While ISGylation is generally linked to tumor progression, emerging evidence suggests that free ISG15 can elicit antitumor immune responses, though the underlying mechanisms remain incompletely defined. Pan-cancer analyses indicated elevated ISG15 expression across all cancer types, with KICH showing reduced expression. Elevated ISG15 in KIRC has been linked to reduced polyubiquitinated proteins, implicating interference with the ubiquitin26S proteasome pathway [32]. Beyond renal cancer, ISG15 is increased in BLCA, UCEC, LUAD, and HNSC irrespective of TP53 status, with even higher levels in TP53-mutant tumors, suggesting roles beyond canonical TP53 signaling. TP53 mutations have been associated with suppressed immune signatures and poor prognosis in HNSC [33], and with tumor-type–specific effects on CD8□ T-cell infiltration [34], although a direct mechanistic link to ISG15 remains unclear. In contrast, ISG15 downregulation in KICH appears independent of TP53 and may reflect its distinct biology, characterized by mTORC1 hyperactivation, PTEN pathway alterations, mitochondrial dysfunction, and oxidative stress [35].

Our study revealed, COAD, KIRC, LGG, and LICH having high ISG15 expression were linked to a lower survival rate [36], whereas in SKCM, MESO and OV [37], high ISG15 expression were associated with an improved survival rate. Our protein-level analysis using CPTAC data corroborated the mRNA expression findings, with the ISG15 protein significantly elevated in all studied cancers except LICH. The observed discrepancy in LICH, where mRNA levels were high but protein levels were low, probably suggests posttranslational regulation and rapid protein degradation mechanisms at play [12,38,39]. Genetic analysis via cBioPortal identified various mutations in the ISG15 gene, including missense mutations, truncating mutations, amplifications, and profound deep deletions. These alterations are significantly associated with shorter PFS, suggesting that genetic modifications of ISG15 can probably contribute to cancer aggressiveness [40]. The presence of both gain-of-function and loss-of-function mutations further emphasizes the context-dependent accompanying cell-and tissue-dependent role of ISG15 in cancer progression [41]. Correlation analysis of the ISG15 genes revealed that IFI35, IF144, OASL, MX1, OAS2, RSAD2, and IRF7 are highly correlated with ISG15 in all 22 cancer types. Reports highlight RSAD2, OAS2, MX1, and ISG15 as significant differentially expressed genes associated with immune function pathways in rheumatoid arthritis [42]. Similarly, Klotz and Gerhauser noted increased expression of the MX protein, OAS1, and ISG15 in response to canine distemper virus infection [43]. The genes encoding ISG15, which are uniquely correlated with a particular cancer, are important in the context of facilitating regulatory mechanisms and functional crosstalk within the tumor microenvironment (TME), which is specific to cancer. One of the uniquely correlated genes, NF-κB, is present in PAAD, and COAD correlation analysis has been reported to be associated with tumor suppression. In myeloma, leukemia, and cervical cancer, ectopic expression of ISG15 and its de-ISGylase USP18 inhibited cell proliferation and induced apoptosis [44]. This tumor-suppressive effect is mediated through disruption of the NF-κB signaling pathway, specifically by downregulating the expression of IKKβ and p65 and decreasing the transcription of the antiapoptotic genes XIAP and Mcl-1[44]. PPI network analysis revealed that ISG15 interacts with key immune regulatory genes, such as STAT1, IFIT1, and IRF7, as well as tumor suppressor genes, such as TP53. UBA7 activated by interferons is known to cause the ubiquitination of ISG15, which facilitates the clustering of many transcription factors that activate the antitumor response and immune evasion in breast cancer [23]. This regulated gene network accompanying the transcription factors STAT1 and STAT2 produces chemokine-receptor ligands for the apoptosis of tumor cells via T cells [45]. The identification of the hub genes STAT1, IFT1, and IRF7 within the network highlights potential pathways for targeted therapeutic interventions. Elucidating the molecular pathways of the hub genes and how ISG15 interacts with key regulatory proteins will help in identifying cancer cell behavior via in vitro and in vivo experiments.

Our analysis also showcased that ISG15 expression is associated with distinct phosphoproteomic states in KIRC and KICH. In KIRC, ISG15 co-varies with a broad phosphorylation landscape involving MAPK, AKT, SRC family kinases, CDKs, and interferon-linked signaling pathways, consistent with an immune-engaged and signaling-adaptive tumor context. Many of these kinase axes have been described as tumor-supportive in specific settings, but their association here likely reflects heightened signaling plasticity and immune interaction rather than direct pro-tumoral activity. In contrast, the ISG15-associated phosphoproteome in KICH is more enriched for cytoskeletal organization, intracellular transport, and stress-responsive kinases, aligning with a structurally maintained and less immune-infiltrated tumor state. In summary, our study reveals a clear divergence in the immune context associated with ISG15 between renal cancer subtypes. In KIRC, elevated ISG15 expression is embedded within a high interferon signaling environment marked by increased immune infiltration, particularly CD8□ T cells and myeloid populations, yet coupled with immune exhaustion signatures, impaired antigen presentation, and unfavorable clinical outcomes. In contrast, KICH displays globally reduced ISG15 expression, preserved methylation, weak coupling to interferon pathways, and an innate-leaning immune association dominated by macrophages and NK cells without coordinated adaptive immune activation. Overall, this study suggests probable context-dependent signaling mechanism within renal cancer subtypes due to different immune signatures, immune infiltration and TME. Additionally, identification of coregulatory genes and PPI networks provide a foundation for future experimental studies aimed at elucidating the mechanistic underpinnings of the role of ISG15 in cancer for the devlopment of targeted therapies.

## 5. Limitations of the Study

Most analyses relied on public datasets, including TCGA, CPTAC, and multiple immunotherapy cohorts, which lack uniform clinical annotations such as detailed treatment regimens, therapy duration, and response criteria. Although extensive survival and prognostic modeling was performed, including multivariable Cox regression and time-dependent ROC analyses, residual confounding cannot be fully excluded. For KICH, limited sample size and fewer survival events may reduce statistical power and contribute to unstable hazard ratio estimates. Discrepancies between ISG15 mRNA and protein expression observed in hepatocellular carcinoma, highlight the limitation of inferring functional protein abundance solely from transcriptomic data. Although CPTAC-based proteomic analyses partially addressed this issue, protein turnover, post-translational modifications, ISGylation dynamics, and degradation pathways were not directly measured and remain speculative in the absence of experimental validation. This study represents correlative associations rather than causative mechanisms, as the analyses were conducted using transcriptomic datasets and computational inference. Functional assays to directly test the role of ISG15 in tumor progression, immune modulation, and epigenetic regulation were not performed. Consequently, the proposed mechanistic interpretations, particularly regarding the divergent roles of ISG15 in KIRC and KICH, should be viewed as hypotheses that require validation through in vitro and in vivo models. Despite these limitations, the integrative, multi-omics approach adopted here provides a comprehensive framework for understanding the context-dependent role of ISG15 across cancers and offers a strong foundation for future mechanistic and translational studies.

## Supporting Information Data

Data 1: Supporting Information Table 1_List of genes similar to ISG15

Data 2: Supporting Information Table 2_Cancer-specific genes correlated with ISG15

Data 3: Supporting Information Table 3_Top 10 hub genes’ rank table

Data 4: Supporting Information Table 4_GO Pathways

Data 5: Supporting Information Figure 1_PPI of ISG15 and its similar genes.

## Competing interest

The authors have no financial or non-financial interest to disclose.

## Conflict of interest

On behalf of all the authors, the corresponding author states that there are no conflicts of interest.

## Author Approvals

All the authors reviewed and approved the manuscript for submission. The work has not been accepted or published elsewhere.

## Author Contributions

S. Chatterjee and J. Mahata contributed equally for joint first author position. Both the authors contributed equally to the research and preparation of the manuscript.

## Funding and Acknowledgements

This research received no external funding. S. Chatterjee is a recipient of a Non-National Eligibility Test (Non-NET) fellowship from the University Grants Commission (UGC), Government of India. J.M. is recipient of the DST-INSPIRE (Department of Science and Technology-Innovation in Science Pursuit for Inspired Research) Post-Graduate fellowship from the Department of Science and Technology (DST), Government of India. We thank the above funding agencies for their financial support.

## Declarations

All data utilized in this study were obtained from publicly accessible repositories, including TCGA, GTEx, CPTAC, and TISCH2. These databases host pre-existing; de-identified patient data collected with appropriate ethical oversight by the respective organizations. Our study did not involve any direct experimentation on human participants or biological specimens. As such, no additional ethical approval or institutional review board (IRB) clearance was required for this work.

## Authors CRediT Statement

**Shrabonti Chatterjee:** Conceptualization, Methodology, Investigation, Validation, Supervision, and Writing - review & editing. **Joydeep Mahata:** Conceptualization, Methodology, Data curation, Formal analysis, Visualization and Writing - original draft.

## Declaration of generative AI and AI-assisted technologies in the manuscript preparation process

During the preparation of this work the author(s) used ChatGPT and Claude opus for improving language and readability. After using these tools, the author(s) reviewed and edited the content as needed and take(s) full responsibility for the content of the published article.

## Notes

### Competing Interest Statement

The authors have declared no competing interest.

### Summary of Updates

Revision Summary We have extensively revised the manuscript in response to the Editors comments and recommendations. The major revisions include: 1. Figures and data presentation: The main figures have been substantially reorganized and simplified to improve clarity and readability. Several supporting panels and complete figures have been moved to the Supplementary Material, while crowded figures have been divided into separate figures to better highlight the key findings. 2. Figure quality: All main and supplementary figures have been prepared and uploaded individually in high-resolution JPEG format to ensure that the figures and labels can be clearly evaluated. 3. Author Contributions: The Author Contributions section has been revised according to the CRediT taxonomy, with each authors specific contributions clearly defined. The contributions in the manuscript have also been aligned with those provided in the submission system. 4. Generative AI statement: The Generative AI statement has been revised according to the journals guidelines. It clarifies the use of AI tools for language editing and readability, while all scientific content, analyses, interpretations, and conclusions were developed and verified by the authors. 5. Figure 1B source: The source of Figure 1B has been clarified in the manuscript and figure legend. The figure was obtained from the GEPIA2 web server, which integrates data from TCGA and GTEx. The source has been appropriately acknowledged. 6. References: The reference list has been reformatted to improve compatibility with the journals reference-checking system. References have been standardized to match the manuscript text formatting, and DOI information has been added where available. Overall, these revisions have substantially improved the figure quality, data presentation, readability, formatting, transparency, and overall presentation of the manuscript.

