## Supplementary Figures V2 for "Integrative pan-cancer analysis suggests ISG15-linked immune and phosphoproteomic divergence between KIRC and KICH"

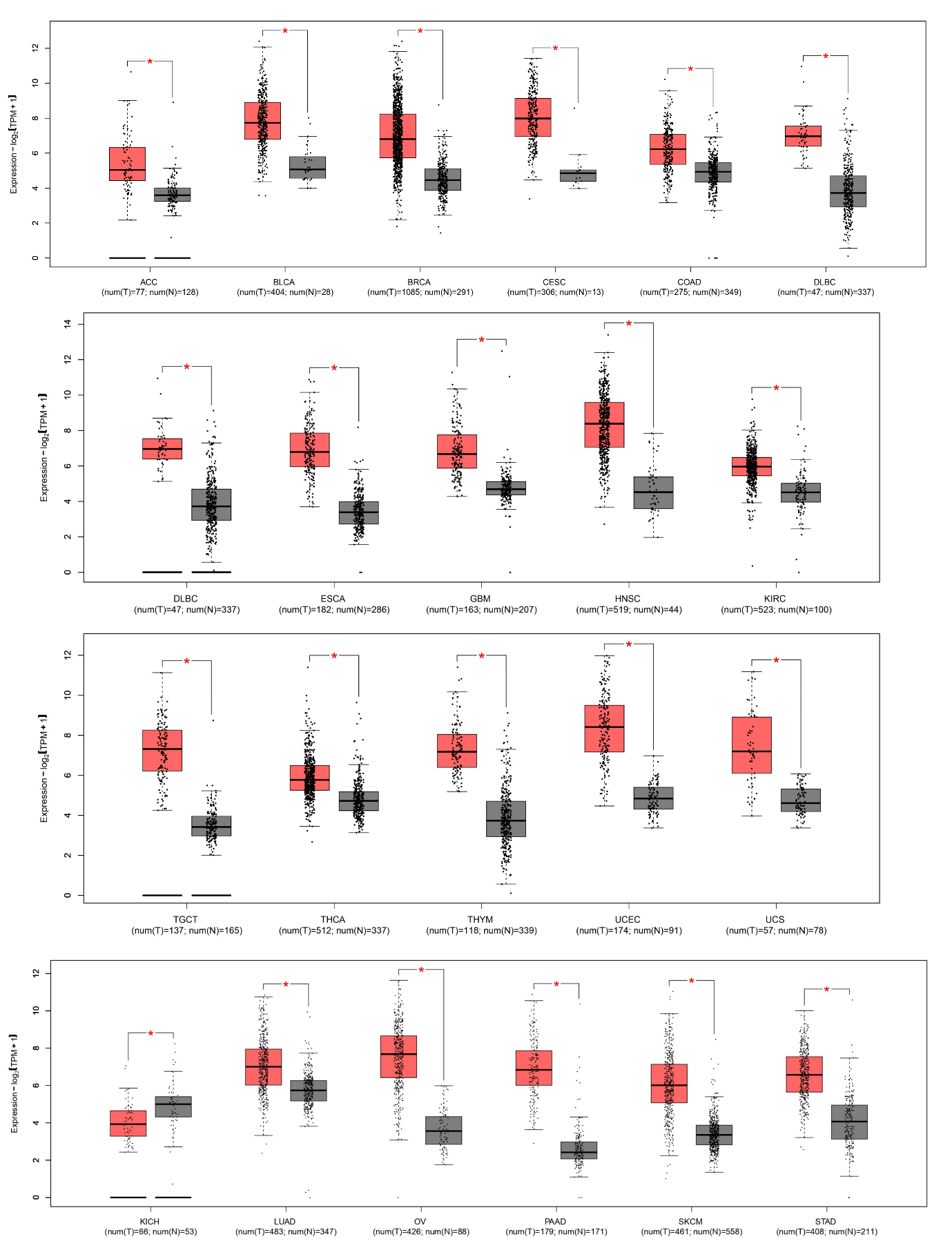

**Supplementary figure 1.** The box plot shows the differential gene expression of ISG15 in 22 TCGA tumors compared with normal samples. The red box denotes tumor sample expression, whereas the gray box reflects expression in normal samples. The horizontal line of each box denotes the mean expression value of ISG15. The X-axis denotes the tumor types with number of samples in Test and Normal. The Y-axis denotes the expression level of ISG15 with log2(TPM + 1). The red star indicates a significant logFC >1.

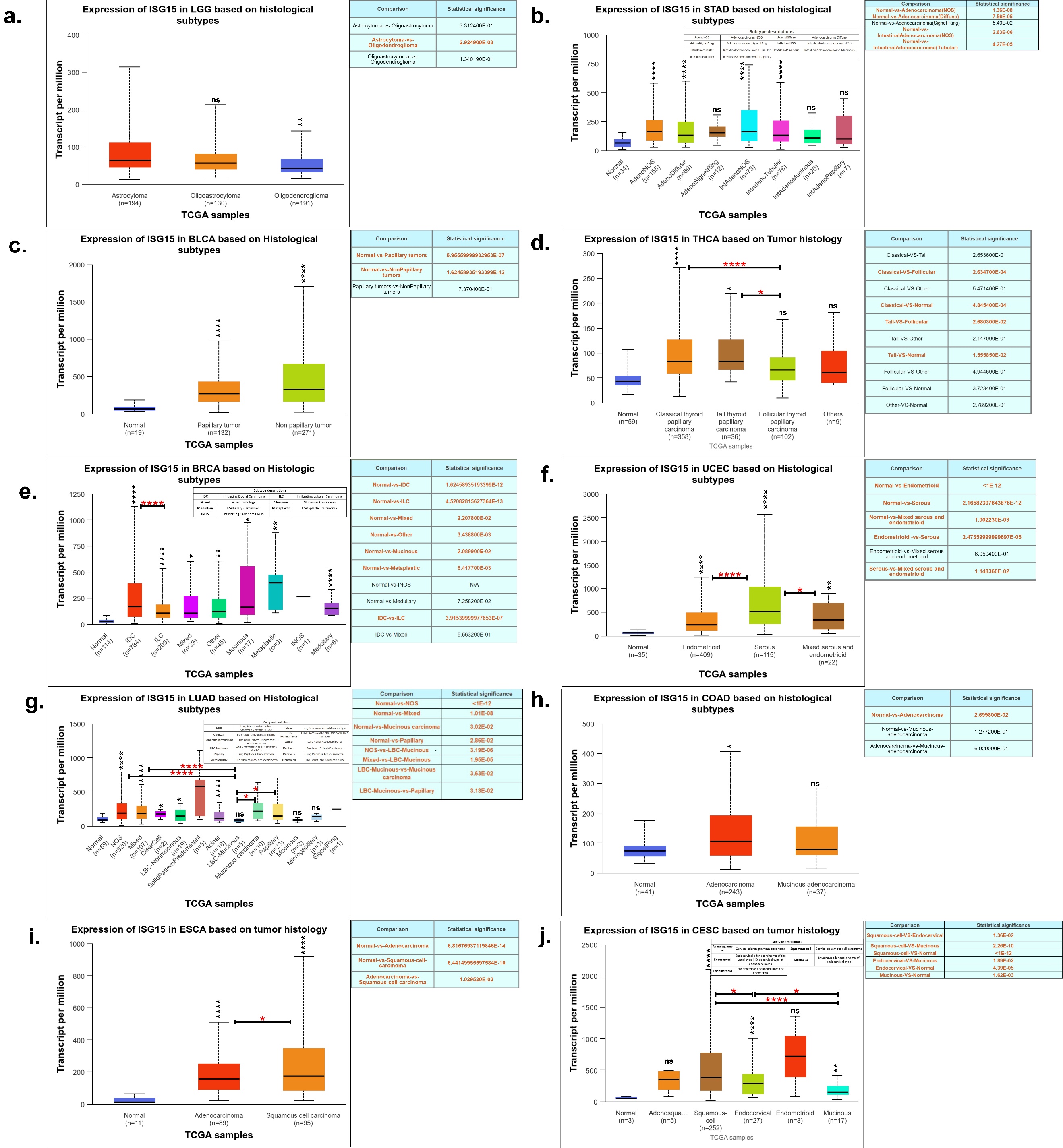

**Supplementary figure 2.** ISG15 mRNA expression expression landscape across TCGA tumors. **(a-j)** Box plot representing ISG15 expression in different cancer subtypes. X-axis denotes the number of samples in different cancer subtype. Y-axis denotes number of transcripts per million in different samples.

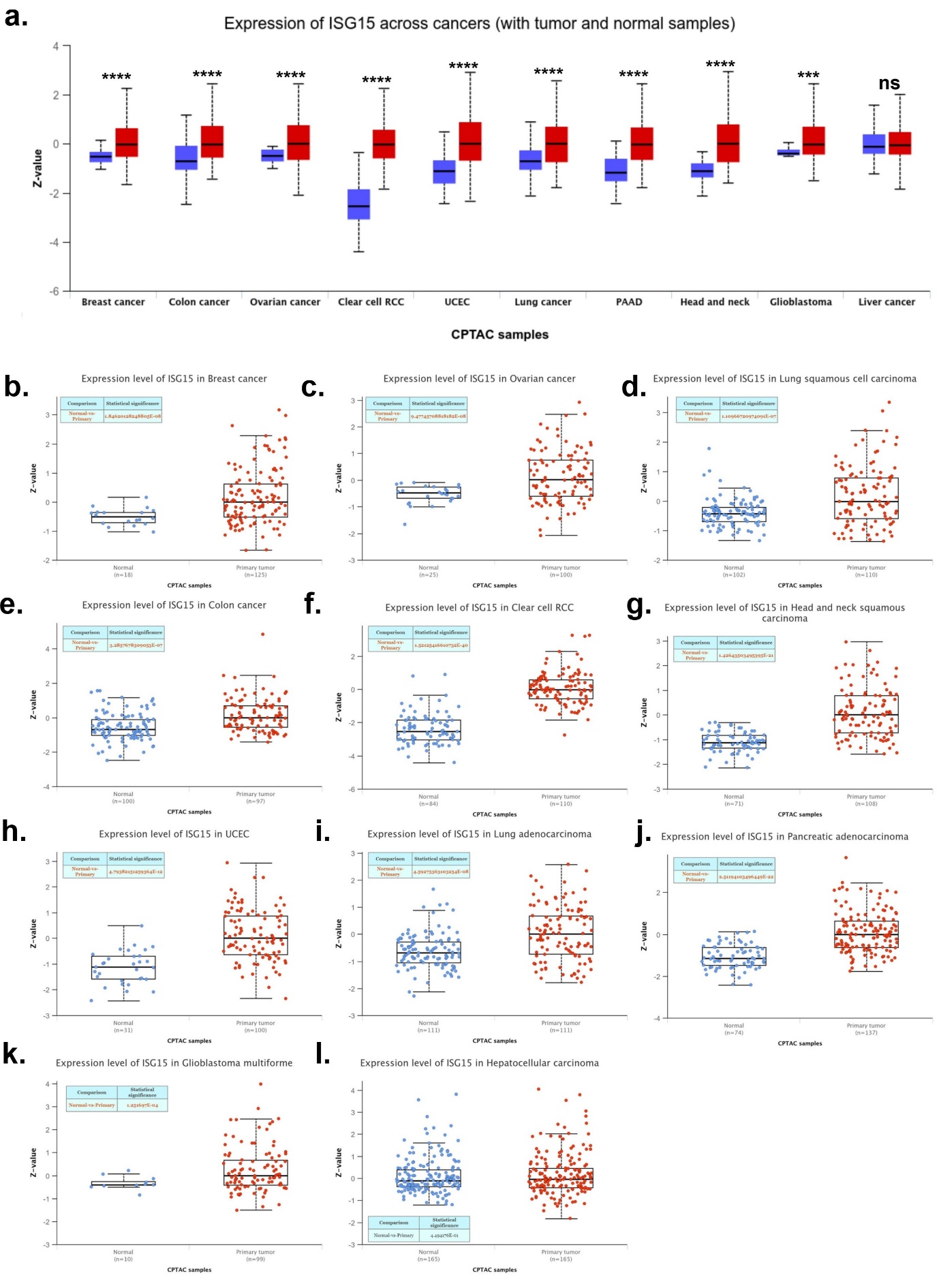

**Supplementary figure 3.** ISG15 protein expression across cancers and stages in CPTAC cohorts. **(a)** Pan-cancer comparison of ISG15 protein levels between tumor and normal tissues. **(b–l)** Cancer type–specific ISG15 expression, shown as normalized Z-scores for tumor and matched normal samples.

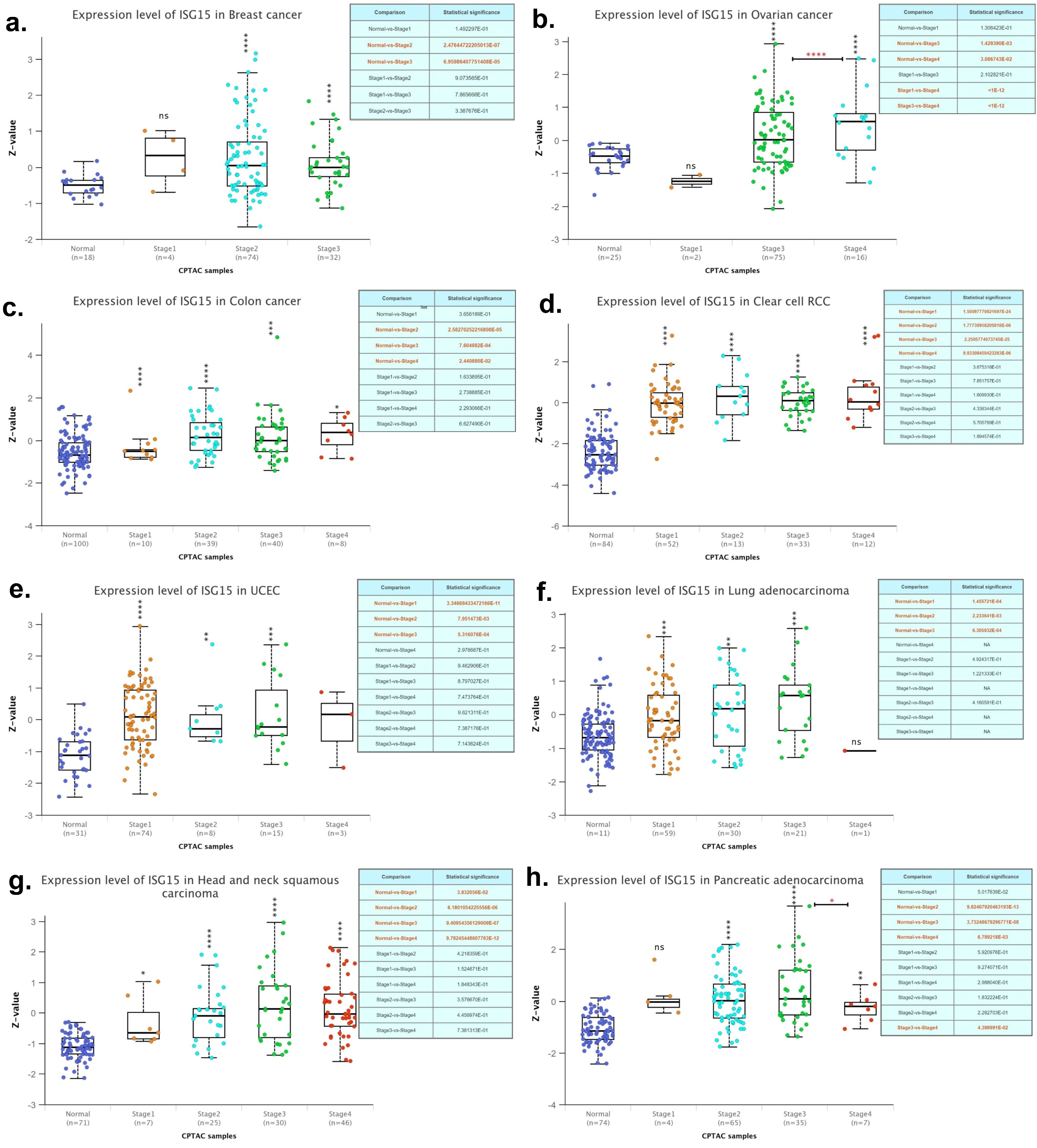

**Supplementary figure 4.** ISG15 protein expression across cancers and stages in CPTAC cohorts. **(a–h)** Stage-wise ISG15 protein expression across cancer types, with stages on the X-axis and normalized Z-scores on the Y-axis. Statistical significance was assessed using non-parametric tests and indicated by asterisks (P < 0.05, P < 0.01, P < 0.001, P < 0.0001); ns denotes not significant.

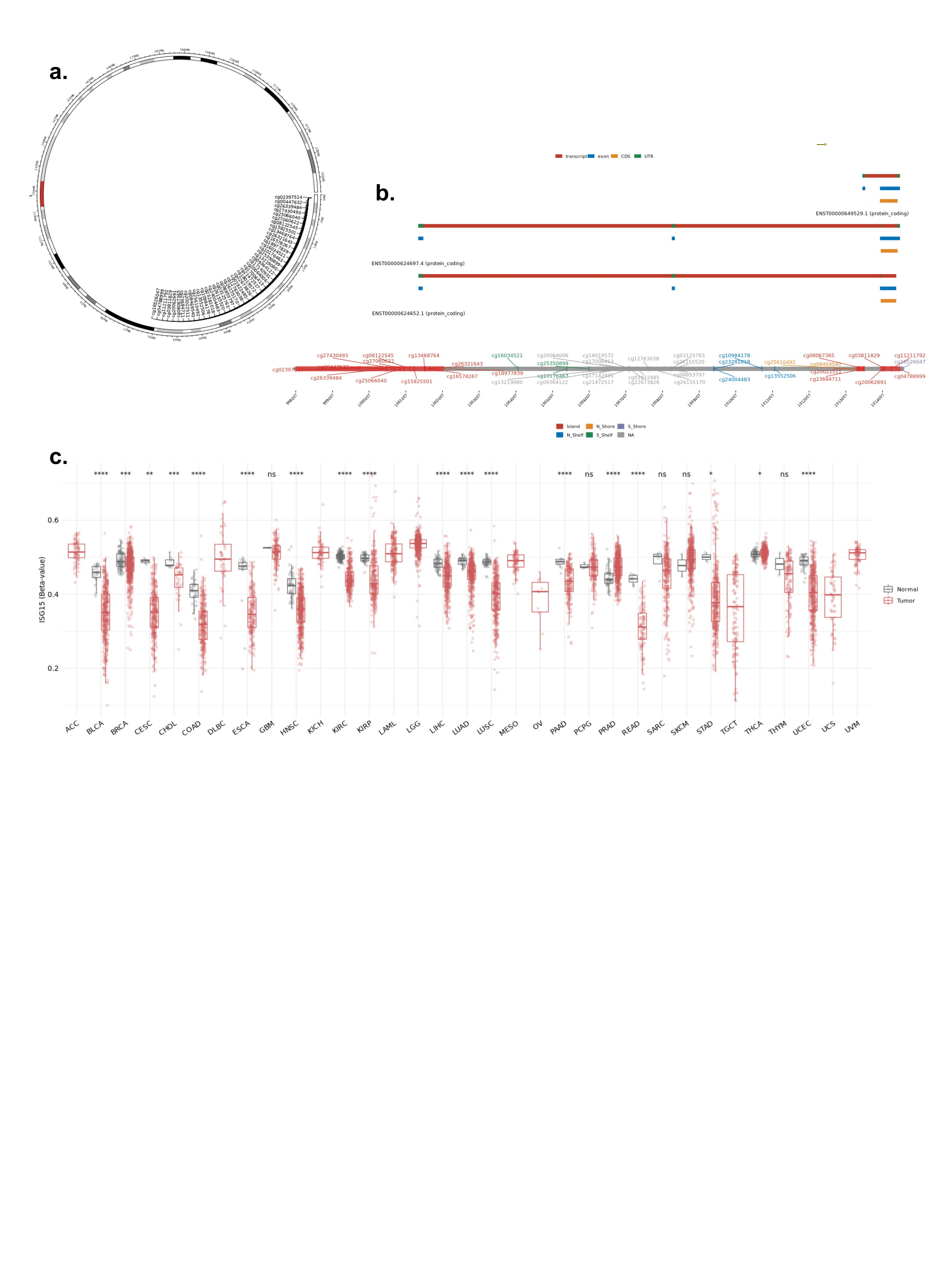

**Supplementary figure 5.** Methylation analysis of ISG15**.** **(a)** Chromosomal distribution of methylation probes associated with ISG15. **(b)** Genomic overview showing 43 methylation sites within the ISG15 gene. **(c)** CpG-aggregated methylation values across all samples.

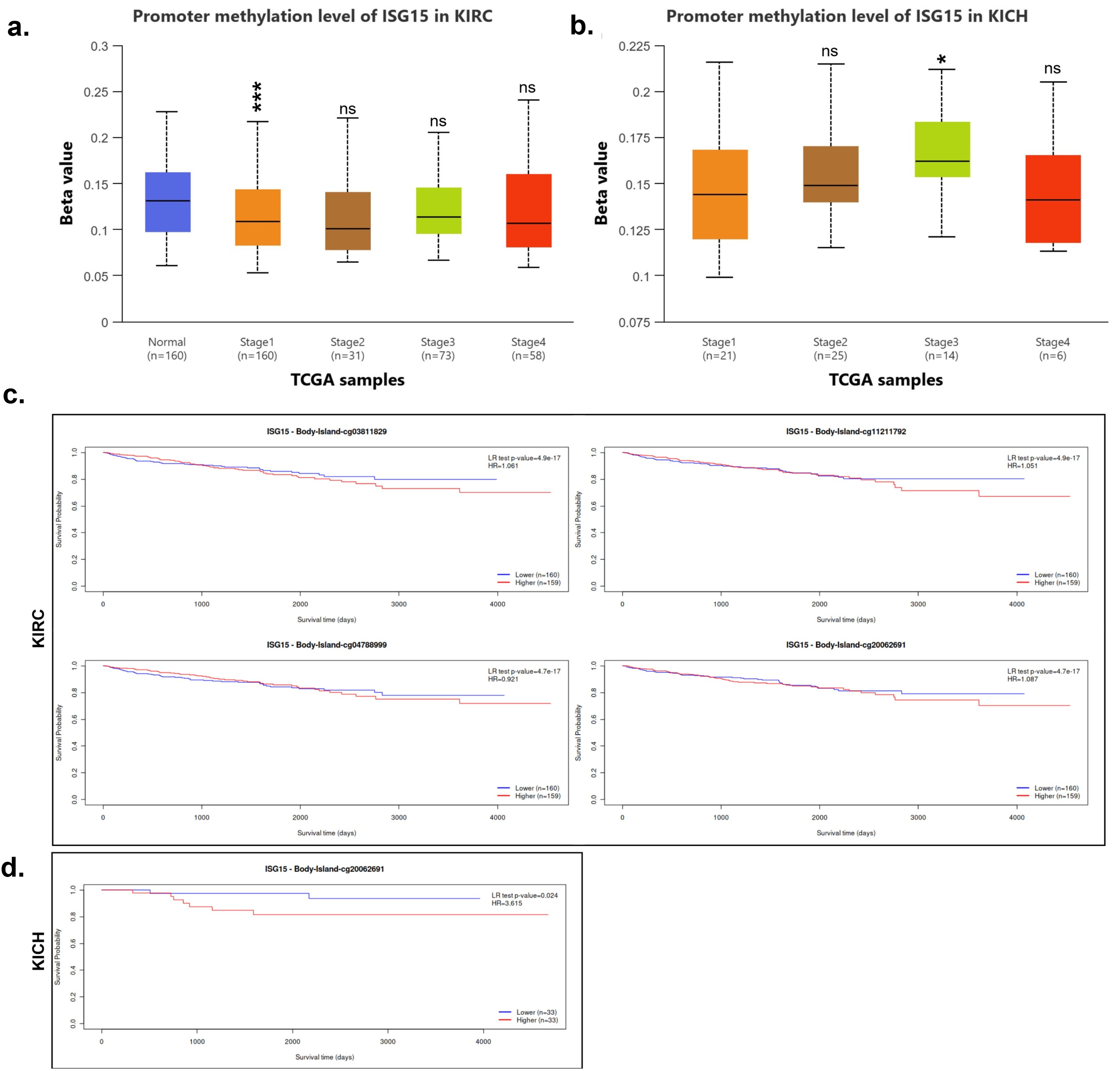

**Supplementary figure 6.** ISG15 promoter methylation and survival associations in renal cancer subtypes. **(a, b)** Boxplots showing ISG15 promoter methylation levels (beta values) across pathological stages in KIRC **(a)** and KICH **(b)** from TCGA; normal tissue is shown for KIRC. Statistical significance across stages is indicated (ns, not significant; * P < 0.05; *** P < 0.001). **(c)** Kaplan–Meier overall survival analyses in KIRC stratified by high versus low methylation at individual ISG15 CpG body island probes. **(d)** Kaplan–Meier overall survival analysis in KICH for an ISG15 CpG body island probe, with log-rank P values and hazard ratios shown, highlighting subtype- and CpG-specific prognostic associations.

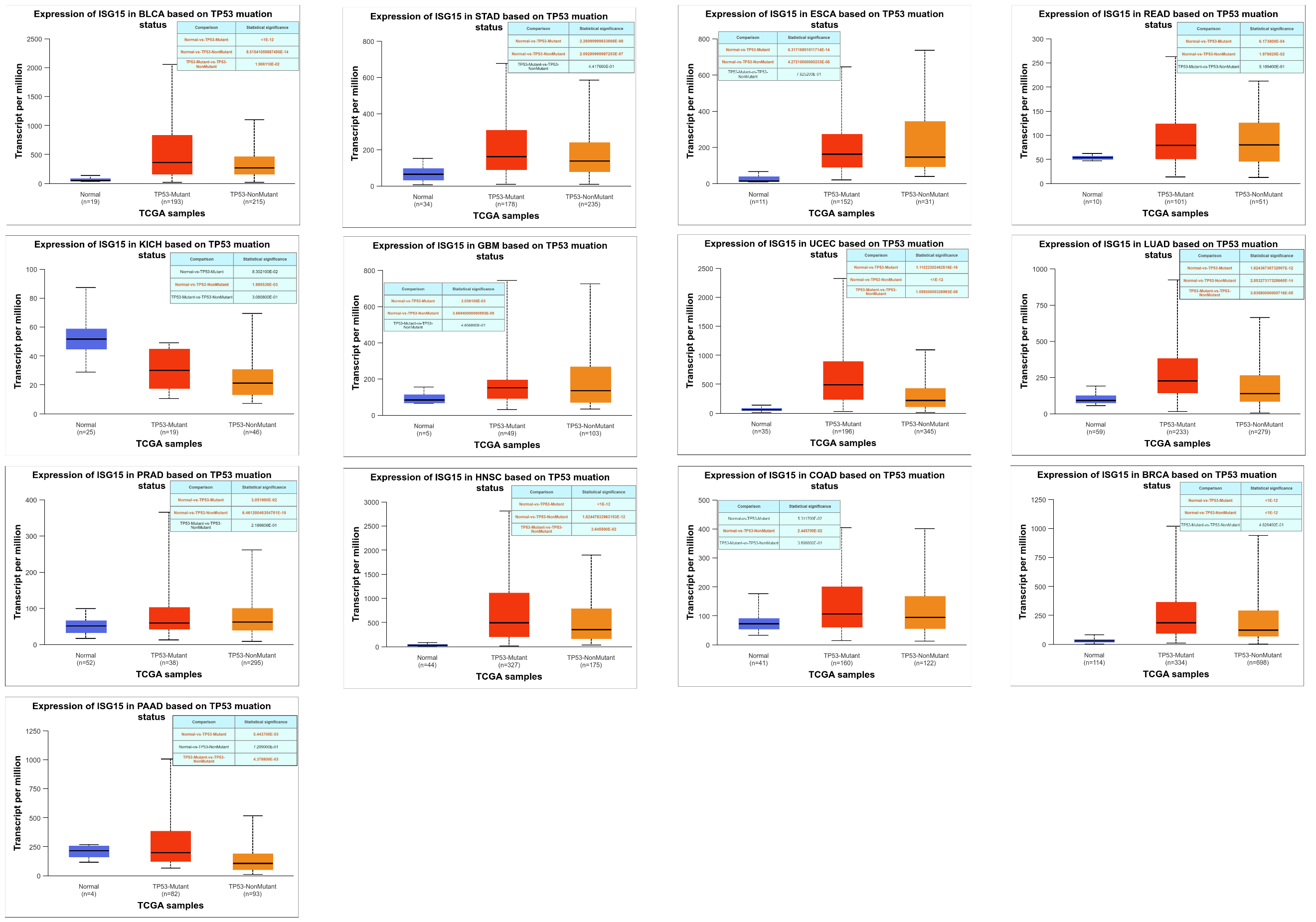

**Supplementary figure 7.** Expression of ISG15 in various cancer types on the basis of TP53 mutation status. Box-plot displaying distribution of ISG15 expression. X-axis represents the different TP53 mutation status whereas Y-axis represents transcripts per million: TP53 mutant tumors (red), TP53 nonmutant tumors (orange), and normal tissues (blue). Each plot compares ISG15 expression across these groups, highlighting variations in expression levels associated with TP53 mutation status. Statistically significant differences between the TP53 mutant, nonmutant, and normal groups are indicated in the respective panels, with p values provided in the top-right inset.

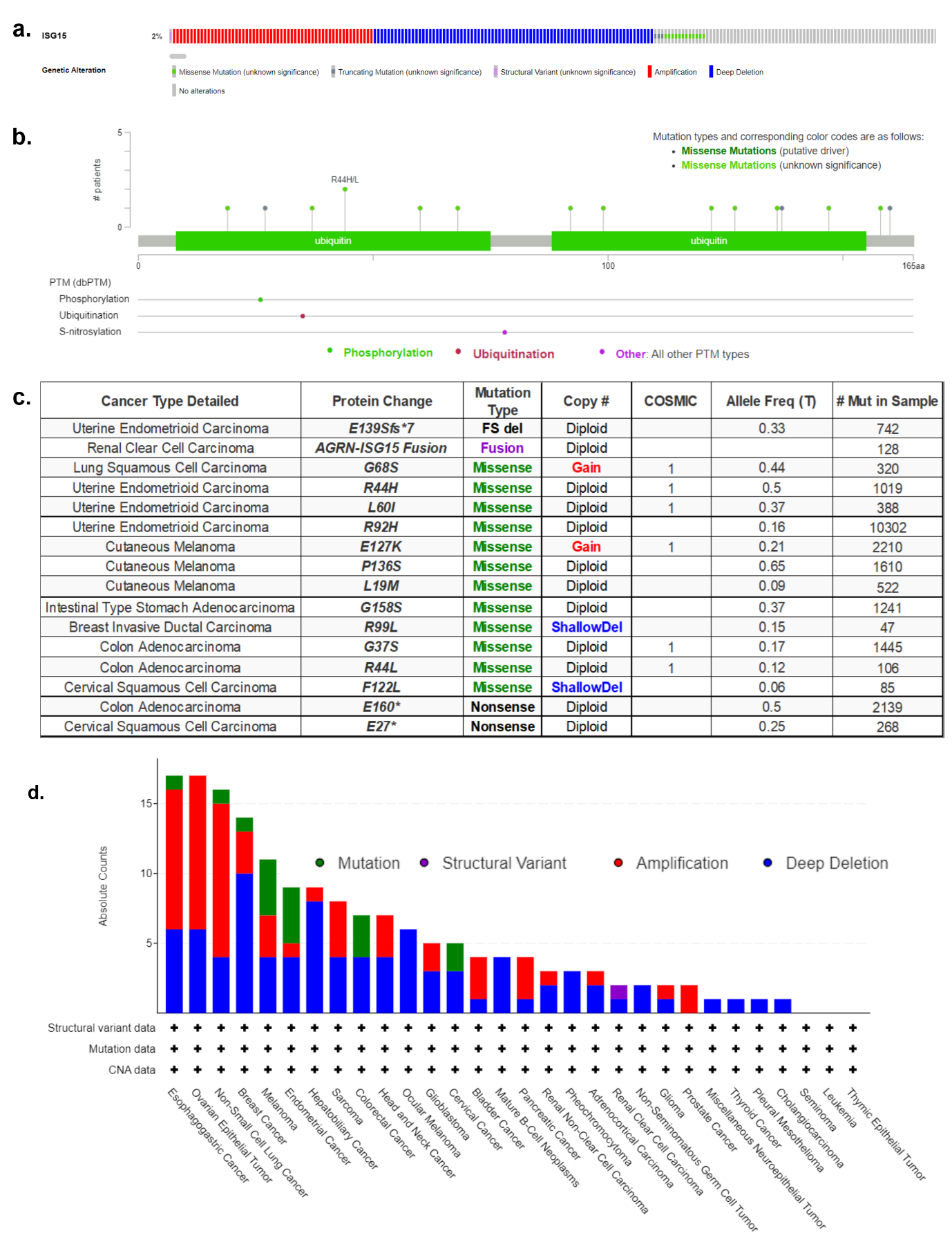

**Supplementary figure 8.** ISG15 genetic alterations in the TCGA Pancancer Atlas. **(a)** Frequency and distribution of ISG15. **(b)** Mutation map of ISG15, highlighting 17 mutations. **(c)** Detailed table listing mutations across cancer types from the TCGA Pancancer Atlas study. **(d)** Pancancer summary of ISG15 gene alterations across various cancer types using data from TCGA Pancancer Atlas studies. Y-axis represents the absolute number of alterations observed, whereas the X-axis lists the different cancer types. Alterations are categorized into four types: mutation (green), structural variant (violet), amplification (red), and deep deletion (blue). Structural variant, mutation, and copy number alteration (CNA) data availability are indicated by the '+' symbols below each cancer type

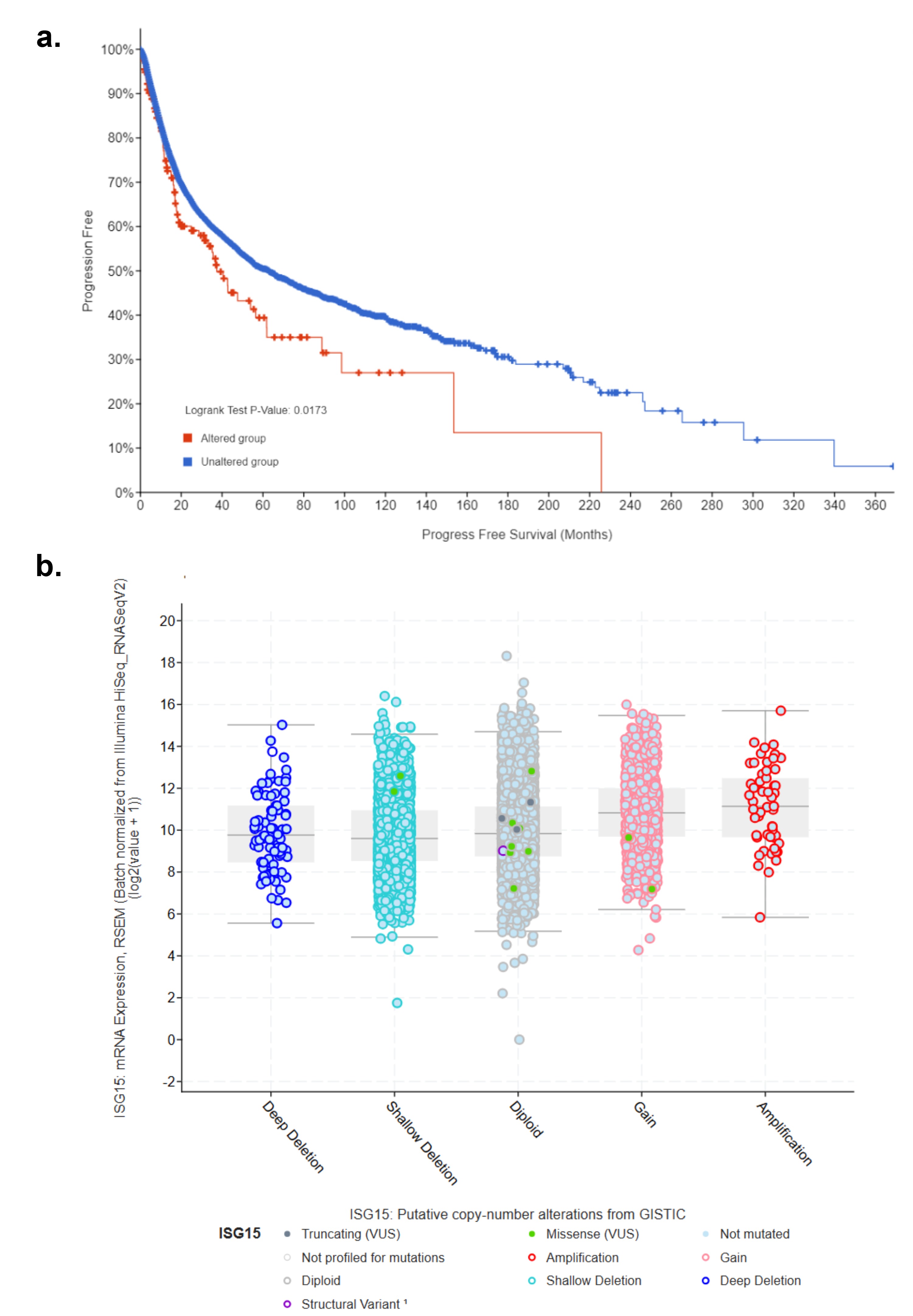

**Supplementary figure 9.** ISG15 gene expression and its impact on cancer prognosis and genomic alterations. **(a)** Kaplan‒Meier survival curve shows the progression-free survival (PFS) of patients grouped by the status of the ISG15 gene. Y-axis represents the percentage of patients who did not experience disease progression, whereas the X-axis represents the time in months. Higher values on the y-axis indicate a greater proportion of patients remaining progression-free. **(b)** ISG15 mRNA expression in relation to copy number alteration (CNA) across different genomic states in cancer. The graph displays ISG15 expression (log2 normalized RSEM values from RNA-Seq data) plotted against five categories of CNAs: deep deletion (blue), shallow deletion (light blue), diploid (gray), gain (pink), and amplification (red)

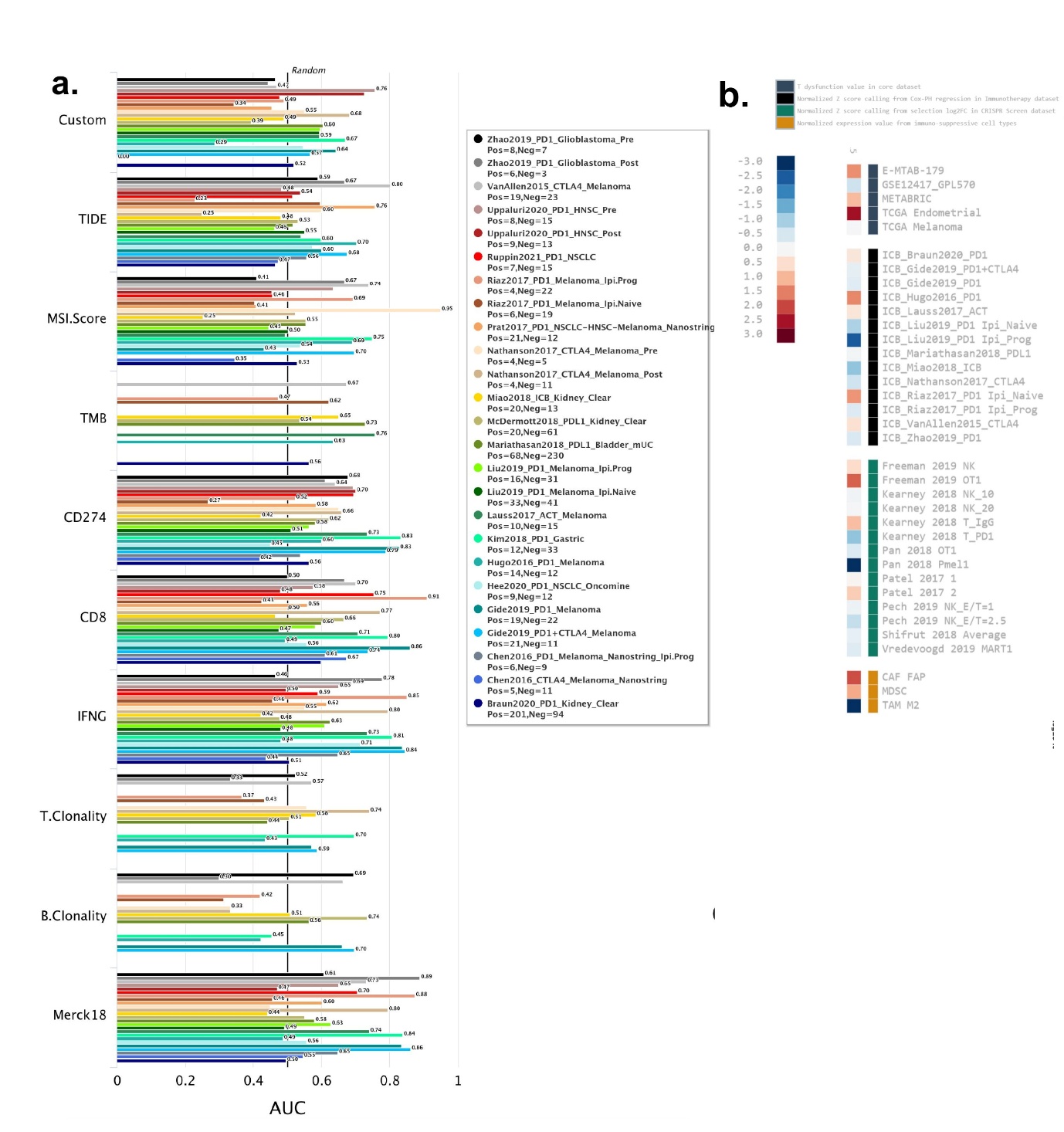

**Supplementary figure 10.** Immunotherapy response evaluation for ISG15. **(a)** Bar plot showing the biomarker relevance of ISG15 (denoted as "Custom") compared to standardized biomarkers in tumor immune evasion across immune checkpoint blockade (ICB) therapy cohorts. **(b)** Immunotherapy response of ISG15 in vitro cell lines after treatment with interferons in cancer cohort. Statistical significance is indicated as *P < 0.05; **P < 0.01; ***P < 0.001.
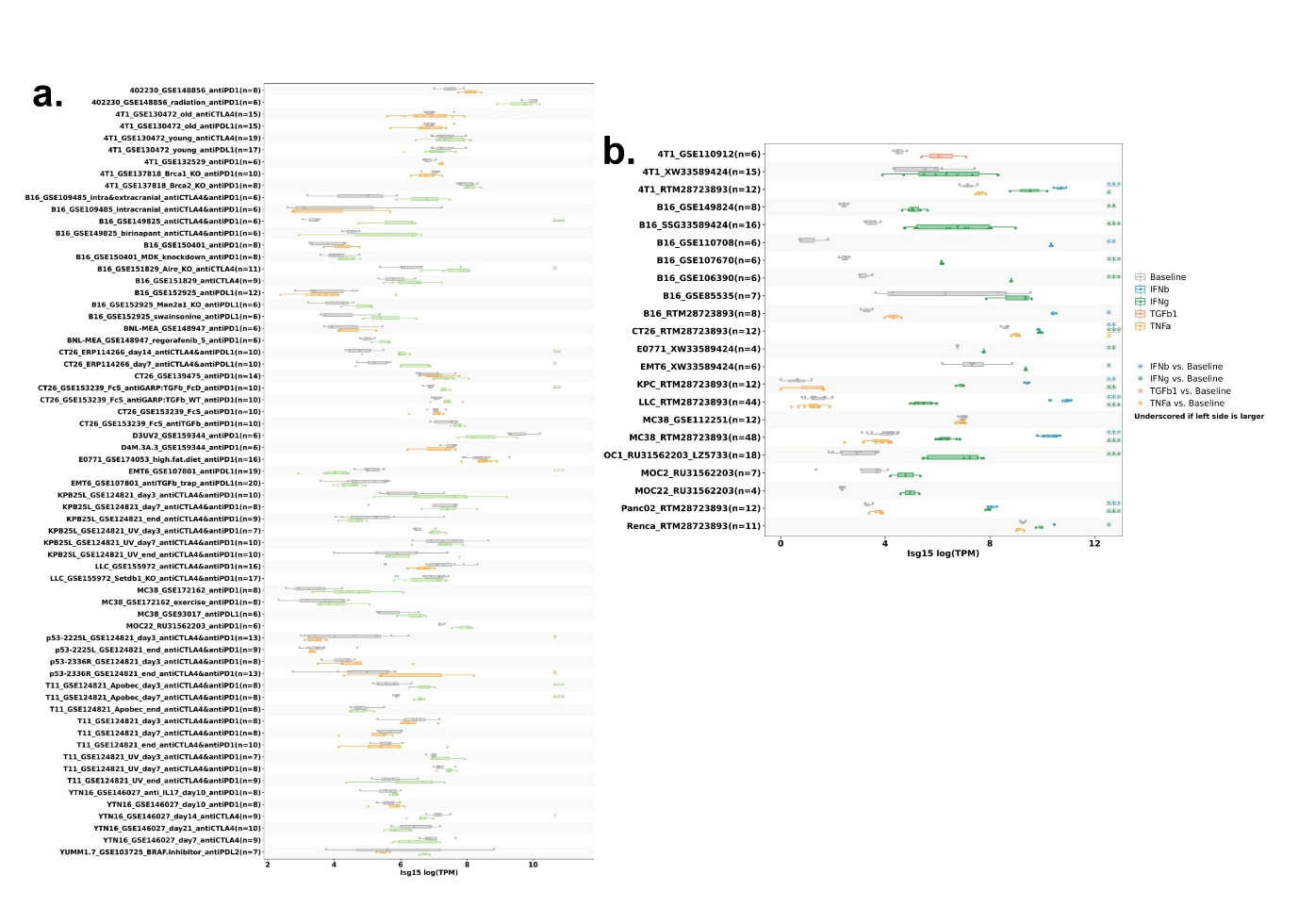

**Supplementary figure 11.** Immunotherapy response evaluation for ISG15. **(a)** Overall survival analysis of ISG15, where the x-axis represents the z-score from Cox proportional hazards regression, and the y-axis represents the -log10 of the P-value. The red line indicates the significance threshold of P < 0.05. **(b)** Immunotherapy response of ISG15 in murein models after ICB treatments in cohort.

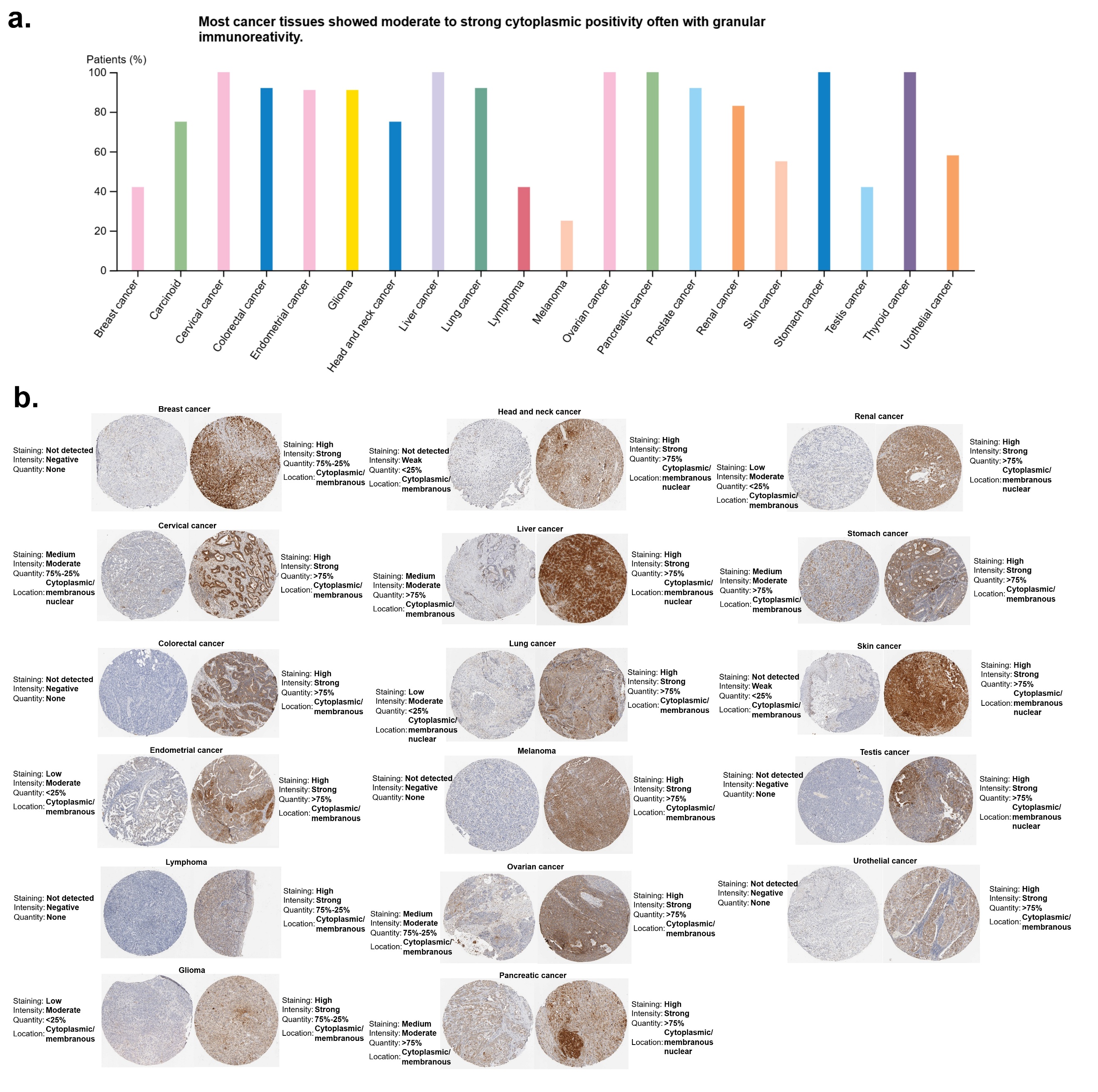

**Supplementary figure 12.** Immunohistochemistry of ISG15. **(a)** Bar diagram showing the percentage of patients with high or moderate ISG15 expression in tumor tissue. **(b)** Immunohistochemistry comparison of ISG15 expression, with no/low/medium staining on the left as control and high staining on the right for tumor tissues.

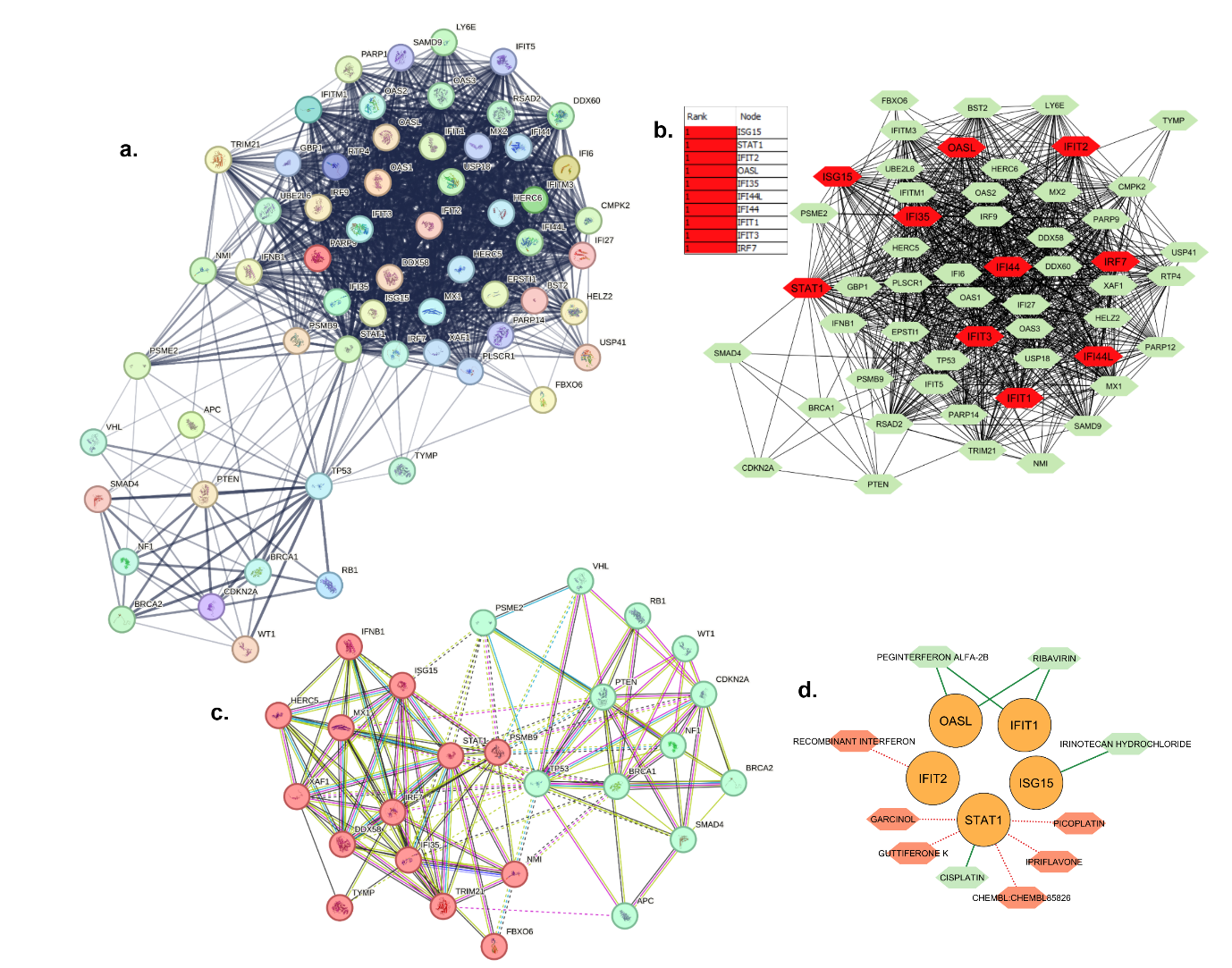

**Supplementary figure 13.** Interaction analysis of ISG15 and similar genes. **(a)** PPI network illustrating the interactions between ISG15, similar genes, and tumor suppressor genes. **(b)** Identification of the top 10 hub genes within the network with its rank table. **(c)** Network diagram showing tumor suppressor genes and their 15 interacting partners derived from ISG15-related genes. **(d)** Drug-hub gene interaction analysis depicting potential drug interactions with the identified hub genes.

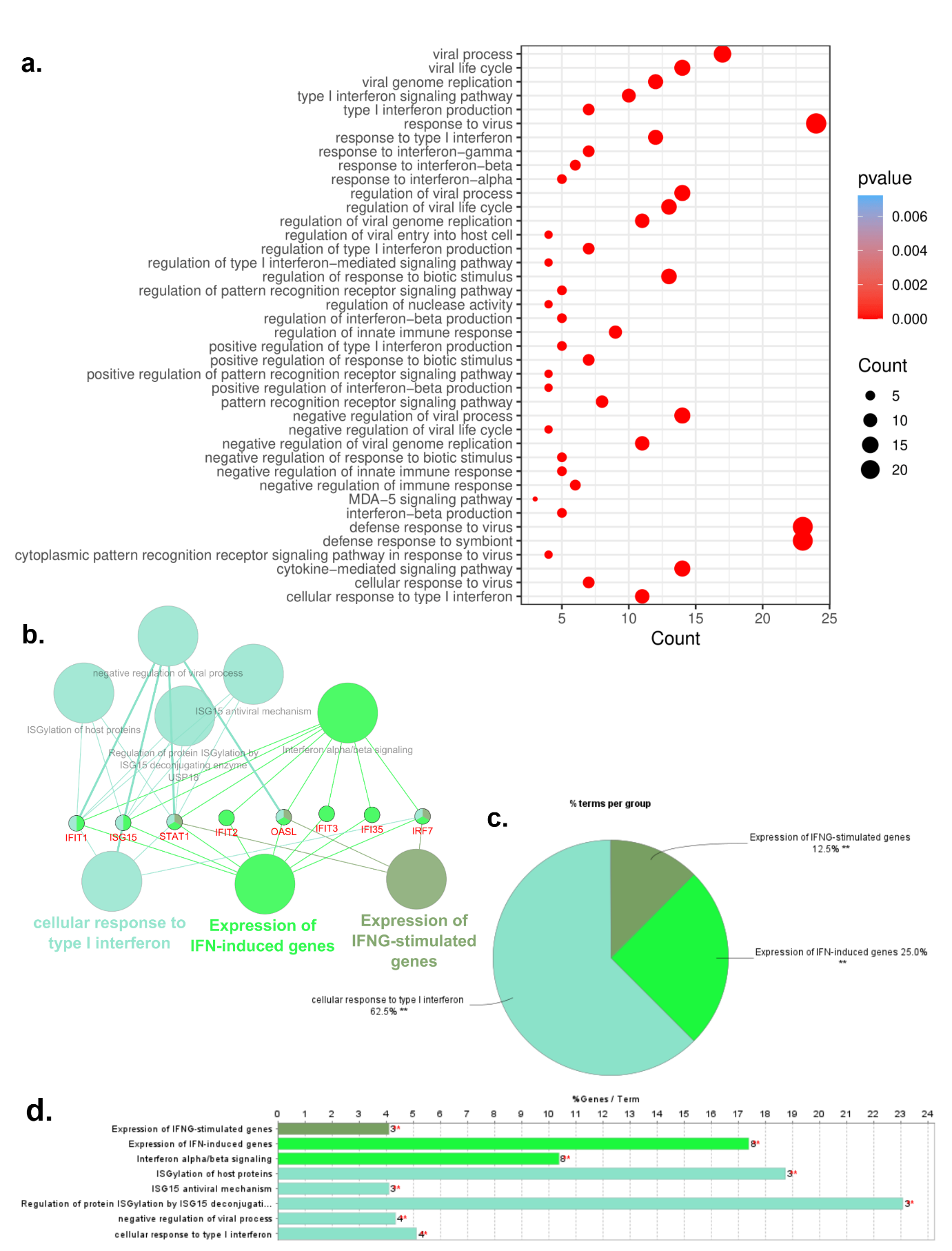

**Supplementary figure 14.** Pathway enrichment analysis of ISG15-related genes and hub genes. **(a)** Dot plot representing the top 40 pathways among 141 significantly enriched GO terms associated with ISG15-similar genes. The dot size indicates the gene ratio, whereas the color gradient (blue to red) represents the lower to higher p value via the Benjamini‒Hochberg method. The X-axis shows the number of genes per GO term, and the Y-axis lists the GO terms. **(b)** ClueGO analysis displaying gene ontology processes, KEGG pathways, immune pathways, and Reactome pathways for the top 10 hub genes. **(c)** Pie chart showing the percentage of terms per group. **(d)** Horizontal bar diagram representing the percentage of genes associated with each term.

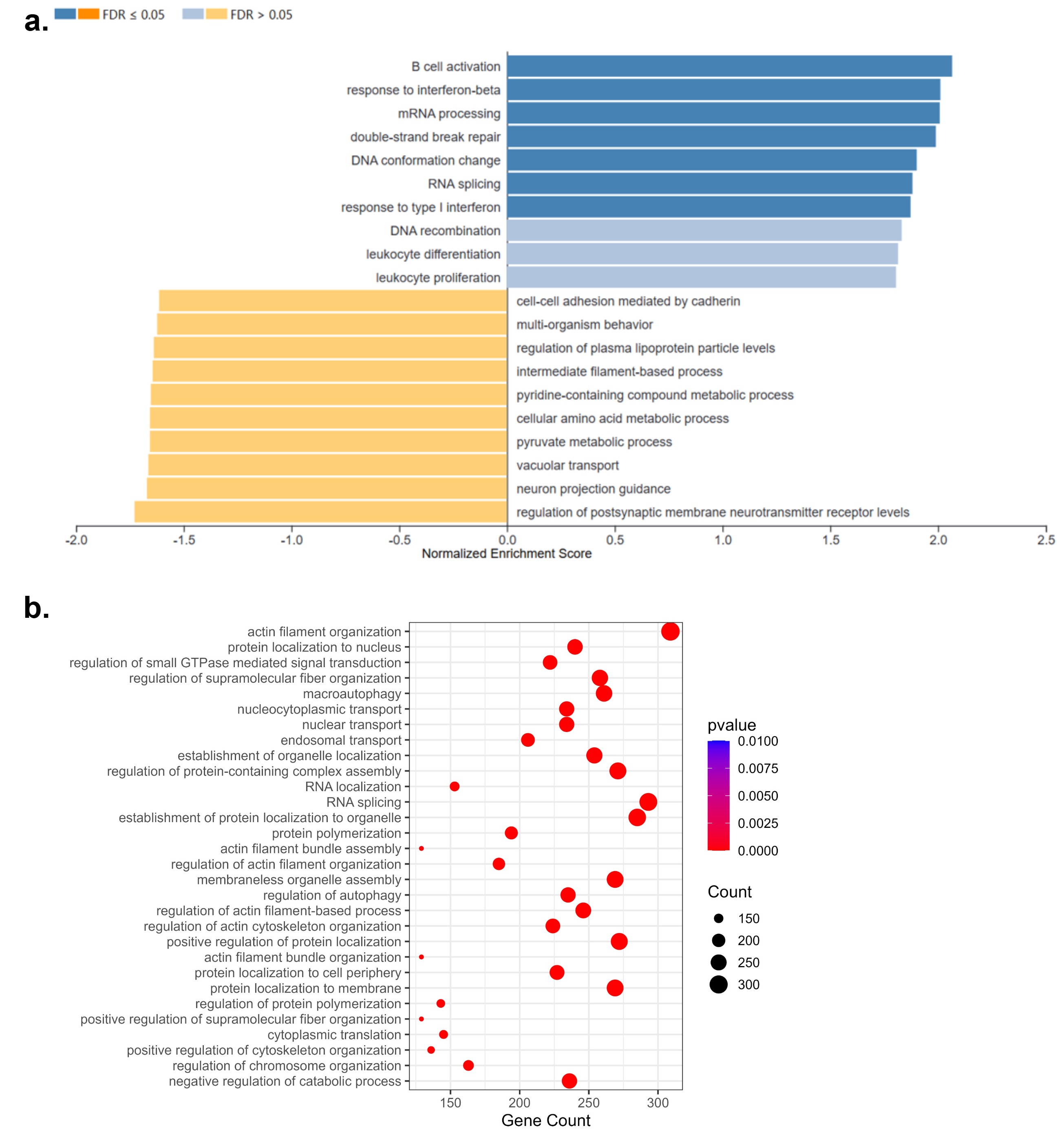

**Supplementary figure 15.** Functional enrichment analysis of ISG15-associated phosphoproteins in renal cell carcinoma. **(a)** GO biological process enrichment of ISG15-associated phosphoproteins in KIRC (FDR ≤ 0.05). **(b)** GO biological process enrichment of ISG15-associated phosphoproteins in KICH based on gene counts and adjusted p values.
